# Pest evolution amplifies projected crop losses under climate change

**DOI:** 10.64898/2026.07.22.740016

**Authors:** Loke von Schmalensee, Camila Rocabado-Peñanco, Elise Cazaux, Lesley T. Lancaster, Greta Bocedi, David Berger

## Abstract

Climate change influences the physiology and population dynamics of ectothermic pests, with major repercussions for global crop production. Yet, how evolution modulates these outcomes remains unclear. We exposed the widespread beetle pest *Callosobruchus maculatus* to 10 years of experimental evolution at different temperatures and quantified thermal responses of life-history traits. Hot- and cold-adapted populations evolved differences in thermal sensitivity, but these were modest relative to evolved differences in trait averages. By leveraging high-resolution temperature time-series we show that the observed evolution translates into cold-adapted genotypes having highest fitness in cold climates and hot-adapted genotypes in warm climates. Hot-adapted beetles maximize fitness in warm climates by increased larval growth, resulting in larger body sizes and higher fecundity. This evolutionary strategy compounds projected crop losses under warming by increasing both intrinsic population growth and per-capita host consumption rates. By year 2100 under intermediate-to-high warming (SSP3-7.0), pest evolution is projected to increase global crop damage potential by +113% from present—twice that expected from warming alone (not accounting for evolution). In major crop-producing areas, where temperatures and the beetles’ host consumption rates are already high, warming increases average crop damage potential by +29%, but evolution amplifies this three-fold to +87%. Evolution also expands *C. maculatus*’ projected colonizable range and in some regions even flips forecasted crop damage reductions into increases. These results identify climate-driven evolution of pest life-histories as an amplifier of agricultural losses and suggest that current projections may significantly understate the threat warming poses to future food security.

## Introduction

Climate change can alter the population dynamics of ectothermic pests via temperature’s influence on metabolism and other physiological processes^1–6^. While outcomes depend on ecological context^7,8^, the general prediction is that climate change will increase crop losses to pests^9^, with major risks to food production worldwide^10^. However, large-scale forecasts of future pest impacts rely on projecting present-day responses into novel environmental settings^6,11^, thus capturing direct, plastic effects of climate but disregarding the potential for evolutionary change in both trait means and plasticity^9,12–14^.

Warming may amplify global pest impacts on agriculture and forestry by expanding the temporal and geographical range of suitable niche space^9,15^, for instance through widening growth seasons^16^, increasing winter survival^17^, or promoting poleward range shifts^18^. Regionally, the effects of warming depend on current temperatures; ectothermic pests in relatively cool temperate regions may enjoy substantial fitness increases while species in hotter regions with temperatures closer to their thermal limits may experience diminished or even detrimental effects of warming^9^, reducing projected impacts on crop yields^6^. Yet, reduced pest impacts due to demographic decline can be counteracted by higher host consumption rates via heat-driven increases in metabolism and pace-of-life^1,6,19,20^, particularly in food-rich crop fields^21–23^.

However, adaptive trait evolution in response to global change can undoubtedly modulate these plastic responses over time (Fig S1)^14,24,25^. Indeed, many pest species exhibit potential for rapid evolutionary responses, mediated by a high demographic turnover that is likely to be further accelerated by warming^21,24,26,27^. This evolutionary potential might be particularly important in already-warm, productive agricultural regions, where demographic declines under further warming are expected to constrain crop losses to pests^6,9^; these declines could be mitigated by thermal adaptation, compounding crop losses beyond current predictions. Indeed, historical data suggest a more positive impact of climate change on crop pest populations than predictions leveraging present-day responses^9^, perhaps precisely because the former inherently accounts for evolutionary processes. Moreover, the specific life-history changes that underlie an adaptive response might carry crucial information, as different trait allocation strategies can result in drastically different host consumption rates even if they yield equivalent fitness outcomes for the pest species in question^4,12,28,29^. Thus far, however, the role of trait-specific evolution remains critically underexplored in quantitative large-scale projections of future pest impacts.

Projecting these eco-evolutionary dynamics requires accounting for the increased frequency, duration, and intensity of extreme temperatures brought on by climate change^30–32^. Ectotherm life-history traits respond to temperature nonlinearly, so even brief exposures to these extremes can have large biological impacts^32–34^, and since diurnal variability constitutes a major source of temperature variation in nature^35,36^, daily extremes play a key role in determining the overall eco-evolutionary effects of temperature (Fig S1)^37,38^. Consequently, even when the intent is to project ectotherm pest responses decades into the future, it is critical to use high-resolution temperature input data with realistic temporal structure in order to make mechanistically grounded predictions^39–41^.

Here, we investigate the extent to which life-history evolution can alter large-scale forecasts of agricultural pest impacts using the seed beetle *Callosobruchus maculatus*, a widespread pest on legumes in tropical and subtropical regions with the ability to rapidly decimate seed storages and cause major crop yield losses^42,43^. We first quantified thermal reaction norms for key life-history processes in multiple genotypes following 10 years of experimental evolution at different temperatures, all conducted on the beetle’s main host, the socioeconomically important grain legume cowpea (*Vigna unguiculata*)^43,44^. We then combined these phenotypic data with globally distributed high-resolution temperature time-series in dynamic life-cycle simulations to study how life-history evolution influences projected crop losses under warming. Under intermediate-to-high warming (SSP3-7.0^45^), evolution doubled the projected global pest impact expected from warming alone, with even greater relative damage increases in regions where cowpea production is highest. Importantly, the projected increases in crop damage potential exceed those expected solely from increased pest population growth, showing that the specific life-history strategies underlying pest adaptation ultimately shape agricultural impacts. Our findings identify life-history evolution in pests as an amplifier of global crop losses in a warmer future, highlighting the importance of accounting for trait-specific evolutionary adaptation when forecasting pest impacts and their associated threats to global food security.

## Results

### Pest trait evolution under different thermal regimes

To determine how climate adaptation shapes pest traits, we quantified evolutionary responses in *C. maculatus* beetles reared on their main host, cowpea (*V. unguiculata*). First, we subjected replicate *C. maculatus* lines sourced from three different origins (Brazil, Yemen, USA) to two novel thermal evolution regimes (cold: 23 °C, hot: 35°C) while keeping founder populations at the benign ancestral temperature of 29 °C (henceforth “ancestors”). We then quantified the evolution of thermal reaction norms by assaying the lines at five temperatures (17/23/29/35/37°C) after 10 years (90–130 generations) of experimental evolution. Evolution under the three different thermal regimes induced repeatable shifts in trait-specific thermal performance across geographic origins (Fig 1). To partition these differences into continuous reaction norms, we employed Bayesian nonlinear models of development rate (day^-1^) and growth rate (g day^-1^) using a thermal performance curve function, and lifetime reproductive success using separate functions for viability (i.e., the probability of producing any adult offspring) and fecundity (conditional on viability; Fig S2).

**Figure 1.**
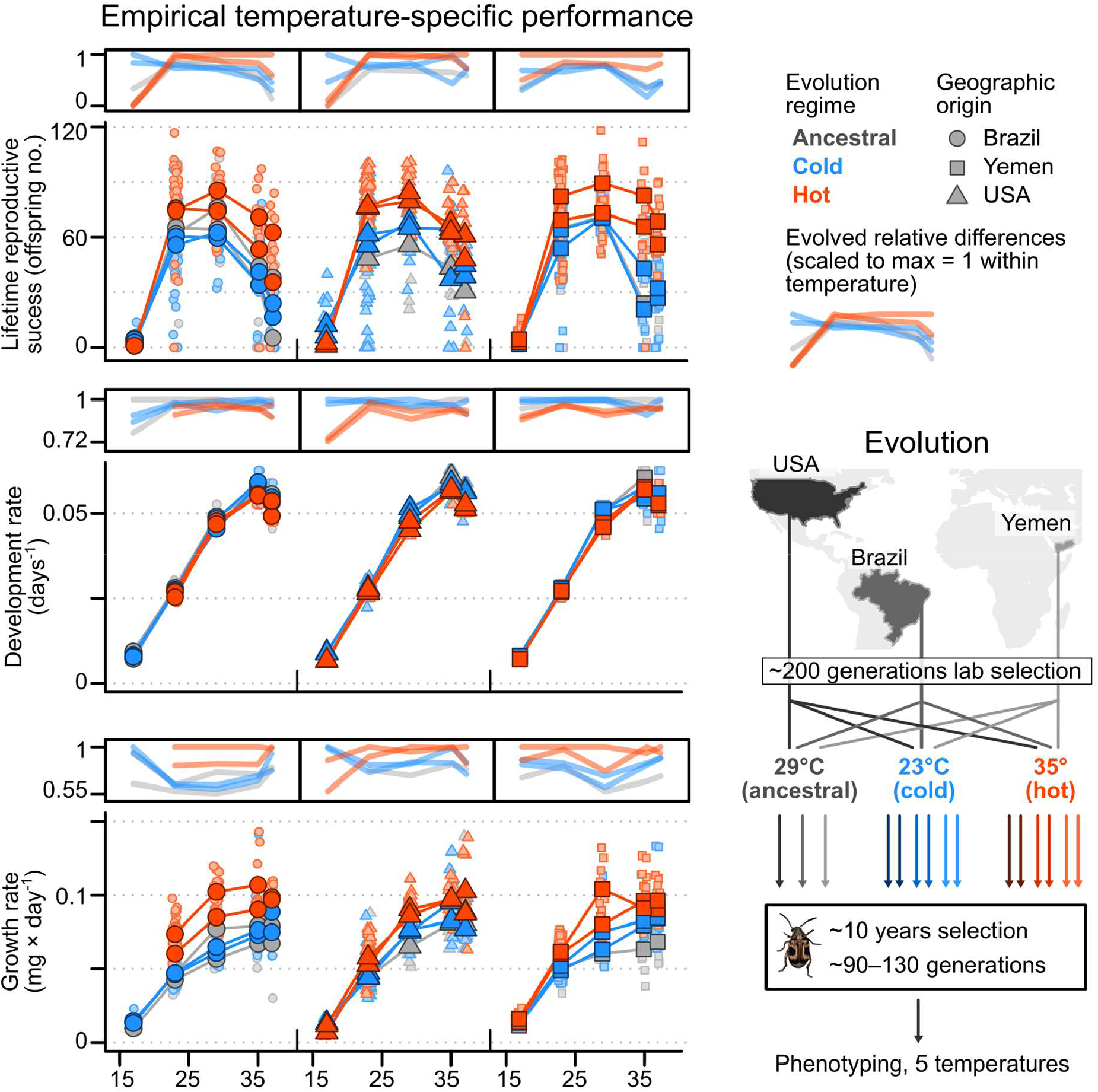
Experimental evolution and evolutionary changes in thermal reaction norms. Populations of *C. maculatus* sourced from three geographic origins (Brazil, Yemen, USA) were maintained under three different thermal regimes (three ancestral founders at 29 °C, six cold populations at 23 °C and six hot populations at 35 °C) for 90–130 generations after an initial ∼200 generations of laboratory adaptation. Populations were then phenotyped across five different temperatures (left panels) to quantify thermal reaction norms; small/light points represent data from individual beetle mating pairs and large/full-colored points represent population averages. Plots above the empirical data show relative differences in trait values, scaled to a maximum of 1 within each temperature. Note that hot-adapted populations of Brazilian origin did not survive at 17°C.

The most prominent observed changes were vertical shifts in performance of the hot-adapted populations, which had evolved larger body size and substantially higher maximal fecundity compared to both ancestral and cold-adapted populations (Fig 2A). As large-bodied genotypes live longer at high temperatures and are generally more fecund (Fig S3)^46–48^, the increase in fecundity in the hot-adapted populations was likely driven by the larger body sizes, resulting from accelerated larval growth but also prolonged development (Fig 2A). Nevertheless, cold-adapted populations had the highest average reproductive output at the coldest temperature (Fig 1), captured in our models by relatively low optimal temperatures for fecundity/viability and relatively wide performance breadth (Fig S4, Tab S1). Moreover, while the cold-adapted populations had a more rapid development than the hot-adapted, the ancestral founders generally developed fastest yet had the slowest maximal larval growth rates (and the smallest adult body sizes; Fig 2A). For detailed parameter estimates and associated uncertainties, see Fig S4 and Tab S1.

**Figure 2.**
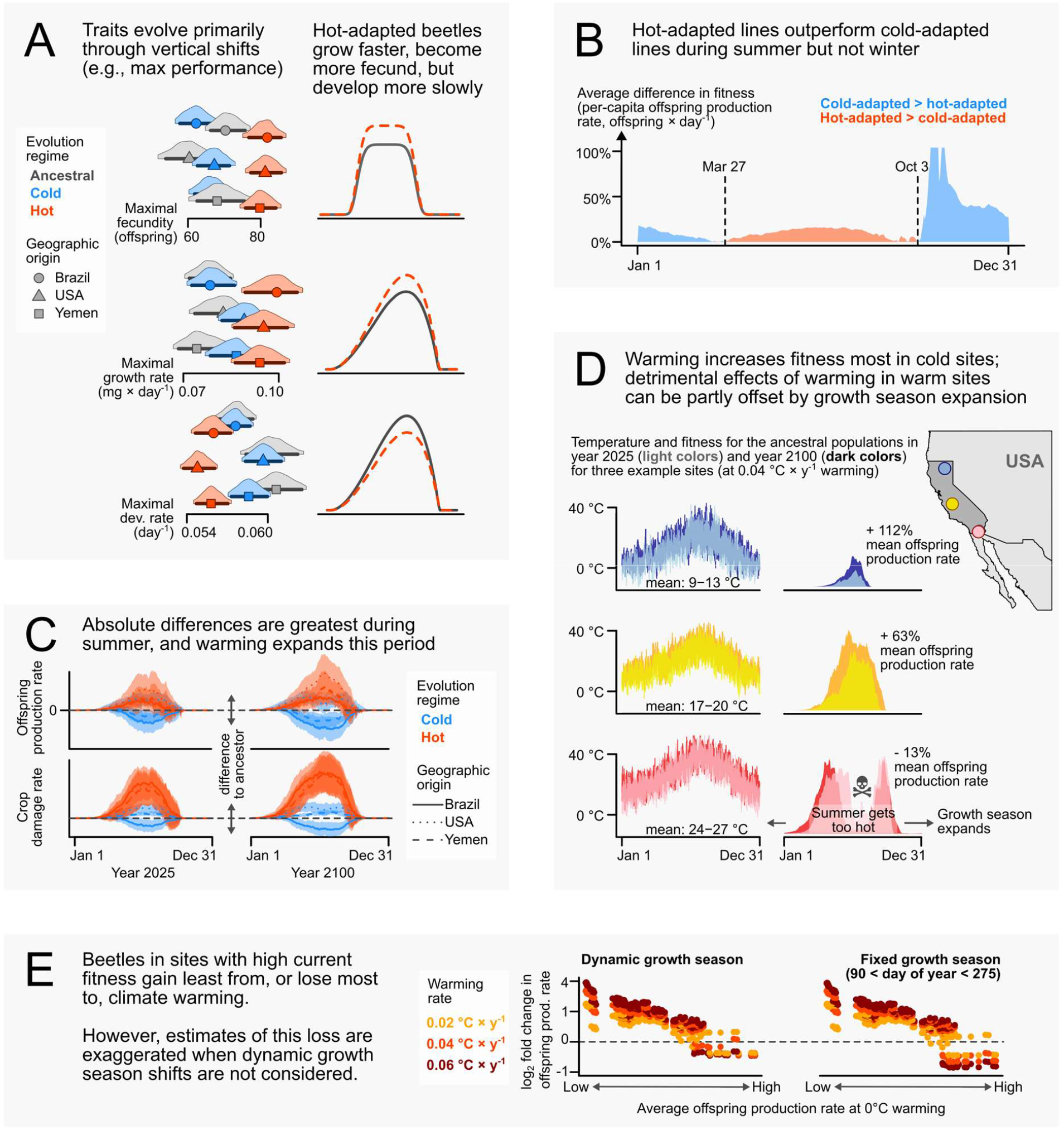
Key insights from subjecting evolved populations to present and future climates. (A) Adaptations to the different thermal regimes were primarily vertical reaction norm shifts; hot-adapted populations have substantially higher maximum reproductive output and growth rates, but slightly lower development rates, compared to ancestors and cold-adapted populations. Densities cover 95% highest posterior density intervals (HPDIs), lines show 85% HPDIs, and points represent posterior modes. (B) Average fitness (per-capita daily offspring production; offspring day^-1^) across sites reveals hot-adapted beetles have higher reproductive output during summer months, whereas cold-adapted beetles perform better during winter. (C) Absolute differences in fitness and crop damage potential (larval growth; g day^-1^) between hot and cold-adapted populations are greatest during summer (plotted as deviations from the ancestors of the same origin; lines show averages, shaded areas show among-site ranges). Warming increases annual fitness and crop damage both via increased maximum rates and by expanding the growth season. (D) Warming increases fitness most at cool sites; at already-warm sites additional warming results in only small fitness increases or even fitness declines due to extreme summer heat. However, such harmful warming can be partly offset by the growth season expanding into spring and fall. (E) Thus, populations with high current fitness gain the least (or lose most) under climate warming. Moreover, disregarding dynamic shifts in the growth season overestimates the detrimental effects of warming on fitness (note: square root-transformed y-axis) which can yield overly conservative estimates of climate-driven crop losses.

Thus, while the evolution of thermal sensitivity in certain traits followed classic predictions under assumptions of antagonistic pleiotropy and trade-offs between performance at different temperatures^20,49^, these differences were relatively modest. Instead, we observed more pronounced evolutionary shifts in the overall performance of specific life-history components, as often observed across latitude in populations adapting to different climates (e.g., “countergradient variation”^50^). This suggests that adaptation during experimental evolution at different temperatures largely proceeded through changes in allocation patterns between different life-history traits (e.g., fast development at the cost of small adult sizes in cold-adapted populations versus slower development but larger adult sizes in heat-adapted populations).

### Thermal adaptation influences seasonal variation in population dynamics and crop damage outcomes

To evaluate how these trait shifts translate into ecological impacts under realistic climate regimes, we derived rate functions from the data (Fig S5) and integrated them over hourly temperature time-series in a dynamic life-cycle simulation (Fig S6)^28^. This simulation allowed us to estimate three key metrics: additive per-capita offspring production (offspring day^-1^), additive crop damage potential via beetle mass production (g day^-1^; Fig S7), and multiplicative population growth potential (geometric mean finite rate of increase, *λ*_geo_). The additive metrics have intuitive units and are better suited for aggregating and decomposing performance, whereas *λ*_geo_ more accurately captures long-term demographic performance and thermal niche suitability. Throughout, we focus on *potential* impacts given the beetles’ thermal physiology and life history: projections were density-independent, isolating intrinsic performance rather than absolute outcomes, which would depend on a multitude of factors that lie beyond our scope.

We explored how seasonal dynamics might modulate the life-cycles of our evolved *C. maculatus* populations across diverse climates, leveraging the fact that California—a major US cowpea producer^51^ and one of our sampling origins—features a highly accessible, high-resolution weather station network over its heterogeneous landscape. We first retrieved hourly temperature data from a diverse set of 20 distinct Californian sites covering a 927 km latitudinal cline. From these data, we then generated site-specific temperature time-series with an hourly resolution and combined them with our empirical thermal reaction norms to run life-cycle simulations under different warming scenarios.

In our simulations, phenotypes of the hot- and cold-adapted populations diverged predictably in performance over the seasons, with hot-adapted beetles generally performing best between mid-spring to mid-fall and cold-adapted lines performing better during colder months (Fig 2B). However, as biological rates drop with decreasing temperatures, average performance is almost entirely dictated by what happens during the warm growing season, which in turn tends to expand with climate warming (Fig 2C). Thus, adapting to cold conditions, where baseline host consumption remains low, generally has negligible impacts on cumulative crop damage potential. Moreover, relative fitness gains from warming are highest in cold sites, as already-warm sites might get dangerously hot during warm seasons. However, such detrimental impact of climate warming on pest fitness can be partly offset by spring and fall temperatures becoming more favorable (Fig 2D,E).

### Trait-specific evolution amplifies effects of warming on crop damage

To quantify how the evolution of individual life-history traits shape crop damage potential under climate warming, we constructed *in silico* phenotypes where a single focal trait (development, growth, viability, or fecundity) was parameterized according to either the hot-adapted or cold-adapted thermal reaction norm while all other traits were constrained to the ancestral response. By comparing these *in silico* phenotypes against both the ancestral baseline and the fully evolved phenotype we partitioned how evolution of each trait contributes to population fitness and crop damage potential under different warming scenarios (Fig 3). The artificial phenotypes were constructed for each geographic origin and thermal evolution regime separately and outcomes were averages across population replicates, resulting in six contrasts (three hot- and three cold-adapted responses) per trait (Fig 3).

**Figure 3.**
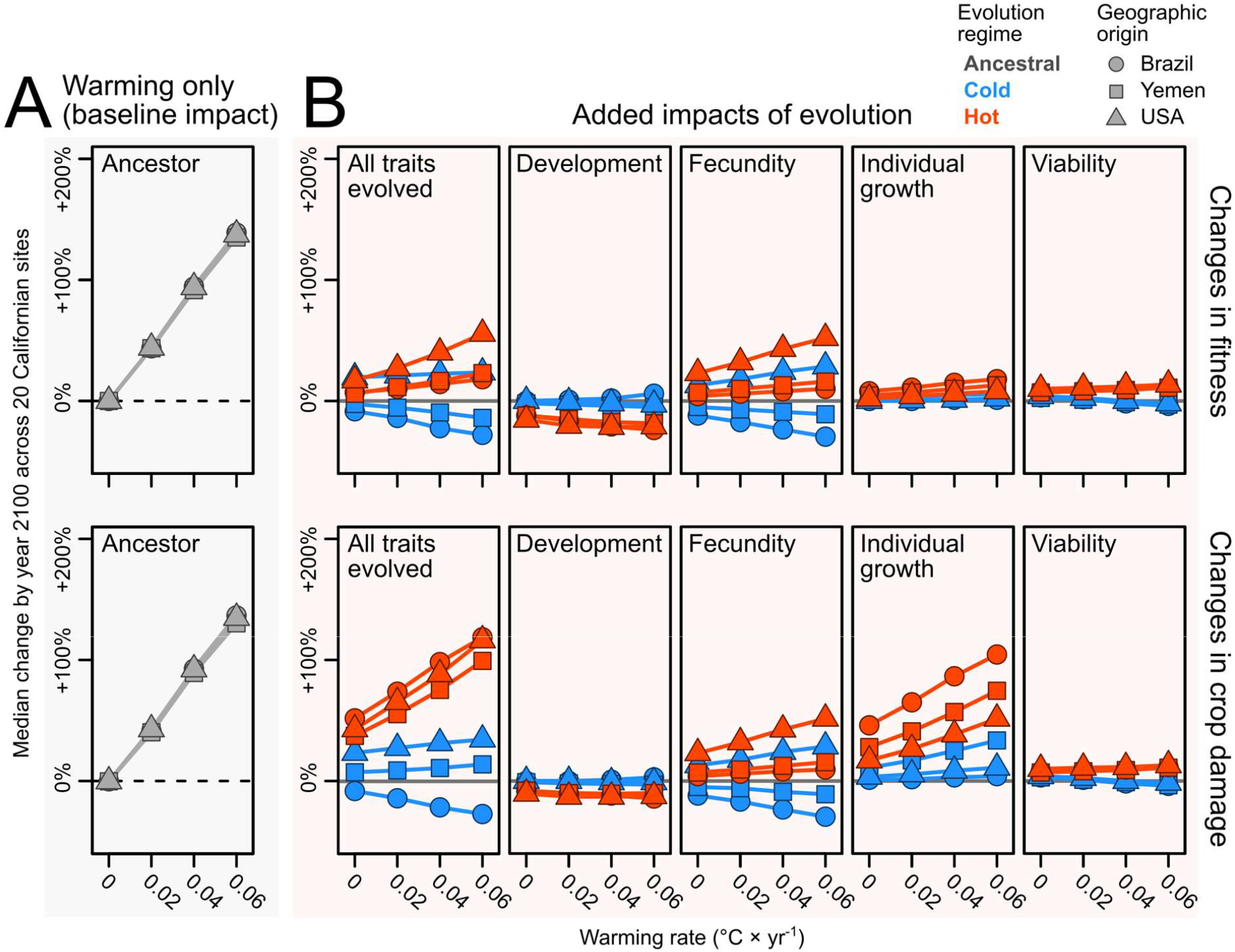
Contributions of trait-specific evolution to pest fitness and crop damage under warming. (A) Median relative changes in fitness (top row; offspring day^-1^) and crop damage (bottom row; g day^-1^) in year 2100 for different warming scenarios compared to present-day levels across 20 Californian sites. (B) Median additional contributions of evolution for the fully evolved genotypes (“all traits evolved”) and four artificial (*in silico*) genotypes in which only a single life-history trait evolves while the other three are fixed to ancestral values, partitioning the ecological impact of thermal adaptation into its underlying trait-specific components. Grey horizontal lines mark zero change from the “warming only” baseline.

Trait-specific responses were different between the two thermal evolution regimes and generally consistent across the different *C. maculatus* source origins within each regime (Fig 3). Compared to present levels at the 20 Californian sites, median baseline warming (ancestral populations, no evolution) affected pest fitness and crop damage potential almost identically: averaged across different warming rates, per-degree warming (°C^-1^) effects were +30% for annualized fitness and +29% for crop damage potential (Fig 3A; Tabs S2,S3). Adding heat-adaptation to these plastic responses amplified both outcomes beyond the warming-only baseline with a substantially larger effect on crop damage than on fitness outcomes. By previous metrics, the full hot-adapted phenotypes added a further +6% °C^-1^to fitness and a further +31% °C^-1^ to the increase in crop damage, relative to the present ancestral state (Fig 3B; Tabs S2,S3).

Dissecting the trait-specific responses reveals that the increased fitness under warming through heat-adaptation is in our system mainly driven by changes in fecundity; the artificial phenotypes with only a hot-adapted fecundity response increased present fitness levels with an additional +5% °C^-1^(*cf*. +6% °C^-1^for the full hot-adapted phenotype; Fig 3B, Tab S2). Interestingly, on its own, the thermal reaction norm for development rate in the hot-adapted phenotypes decreased fitness with warming, indicative of performance trade-ofs across traits.

Yet, in the hot-adapted lines, fitness increases only partly explained increases in crop damage potential; evolving fecundity alone increased crop damage potential by +4% °C^-1^ (*cf*. +31% °C^-1^ for the full hot-adapted phenotype; Fig 3B, Tab S3). Instead, individual growth rates contributed most to the amplified crop damage potential seen in the hot-adapted phenotypes at +20% °C^-1^ (Fig 3B, Tab S3). Cold-adapted phenotypes generally had lower fitness as temperatures rose from present levels (−3% °C^-1^) but they still displayed a slight increase in crop damage potential (+4% °C^-1^) owing to their somewhat higher growth rates than the ancestral phenotypes (Fig 3B, Fig S4, Tabs S1–S3). The conclusions remained qualitatively unchanged following sensitivity analyses (*Methods*, Tabs S4,S5).

### Evolution under constant temperatures translates into geographically structured local adaptation

To forecast the potential global footprint of pest evolution in *C. maculatus* and investigate how our fine-grained insights (Figs 1–3) scale up to patterns across major agricultural production zones worldwide, we re-fitted “global” reaction norms for each selection regime by estimating evolved responses jointly across the three geographic sampling origins (Fig 4A; Tab S1). This pooling also improves computational tractability and is supported by the general consistency of responses within selection regimes (Figs 1,S4,S5; Tab S1). We used these global models to run our dynamic life-cycle simulation across all land between latitudes 60 °S and 60 °N at 1° × 1° spatial resolution with full annual cycles of hourly temperatures, both under present-day conditions (ERA5-Land reanalysis)^52^ and under a spatially and seasonally resolved intermediate-to-high climate warming scenario (WorldClim v2.1, SSP3-7.0, years 2081–2100)^53^.

**Figure 4.**
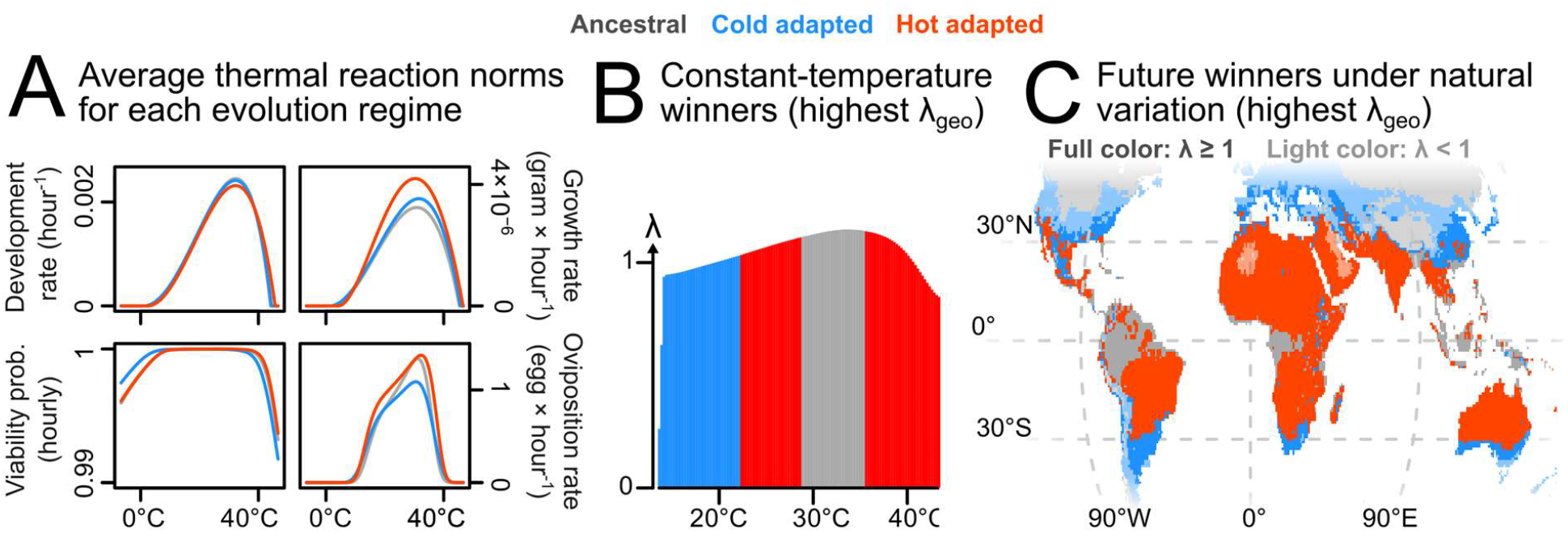
Subjecting evolved populations to regional temperature regimes reveals patterns of local adaptation. (A) Estimates (based on posterior modes) of the four life-history processes (development, growth, fecundity, viability) for the evolution regimes, estimated jointly across the three geographic sampling origins. (B) Predicted geometric finite rate of increase, *λ*_geo_, as a function of constant temperature. Bar colours denote the evolution regime achieving highest *λ*_geo_at each temperature, reflecting adaptation to their respective selective environments. (C) Evolutionary winners, defined by the highest *λ*_geo_, in a global projection at a 1° × 1° spatial and hourly temporal resolution under a spatially and seasonally resolved intermediate-to-high climate warming scenario (SSP3-7.0, years 2081–2100).

While suitable for the seasonal and trait-specific analyses outlined above, our linear offspring production rate (offspring day^-1^) does not account for the compounding effects of demographic growth, potentially misrepresenting long-term persistence. In our global simulations, we therefore used the geometric mean finite rate of increase (*λ*_geo_) over a whole year to determine long-term demographic growth potential, calculated by combining viability, development times, and oviposition schedules in an approximation of the Euler-Lotka equation. Values of *λ*_geo_ below one indicate population decline and eventual local extinction while values above one indicate thermal niche suitability.

Running the life-cycle simulation for the three evolution regimes over a range of constant temperatures revealed clear and predictable patterns of specialization driven by evolutionary thermal history (Fig 4B). The cold-adapted phenotype achieves the highest fitness (*λ*_geo_) at cold temperatures whereas the hot-adapted phenotype does best under high heat. At intermediate temperatures including the ancestral rearing conditions of 29 °C, the ancestral phenotype wins. Additionally, globally upscaling these responses under climate warming using naturally variable high-resolution temperature data yields distinct geographic zones of regime-specific dominance (Fig 4C). The cold-adapted beetles have highest population growth potential primarily in high-latitude and high-altitude regions with frequently low temperatures. Beetles from the ancestral thermal regime dominate a narrow band around the equator with high mean temperatures and buffered thermal variability (matching their evolution regime temperature), whereas hot-adapted beetles have highest population growth potential across the vast majority of the tropical and subtropical regions, where thermal regimes frequently encompass more extreme heat.

### Pest adaptation amplifies projected crop losses across major agricultural producers

To translate the patterns of local adaptation into global projections of agricultural risk, we calculated year-averaged crop damage potential across the global grid. For each 1° × 1° grid cell, we assigned a damage potential corresponding to that of the phenotype with the highest fitness (*λ*_*geo*_) in that environment, setting the projected crop damage potential to zero in regions where *λ*_*geo*_< 1; that is, we only included thermally habitable grid cells. We also retrieved globally distributed coordinates from the Genesys database^54^ for accessions of cowpea (*V. unguiculata*). We then filtered the accession data for farmed/cultivated habitats in main cowpea producing countries (as identified by Boukar *et al*. [2019]^44^ with the addition of India; see *Methods*).

Under present-day conditions, the highest baseline crop damage potential is concentrated across tropical and subtropical environments, including the main cowpea production zone of sub-Saharan west Africa (Fig 5A). Under climate change by year 2100, direct effects of warming alone (excluding evolution) are projected to expand *C. maculatus* suitable thermal niche space polewards (*λ*_geo_ goes ≥ 1) while causing local extinctions (*λ*_geo_ falls < 1) mainly around hot and arid regions such as Sahara/Sahel and the Arabian peninsula. These projected extinctions represent a decrease in fitness due to extreme heat, which drives projected crop damage potential down also in surrounding regions, even when it does not reduce *λ*_geo_ below 1. In general, however, climate warming increases the crop damage potential of the ancestral phenotype by an area-weighted average of +58% across all grid cells (Tab 1A), marking a substantial upward shift mainly driven by an accelerated population turnover rather than range expansions (Tab 1B).

**Figure 5.**
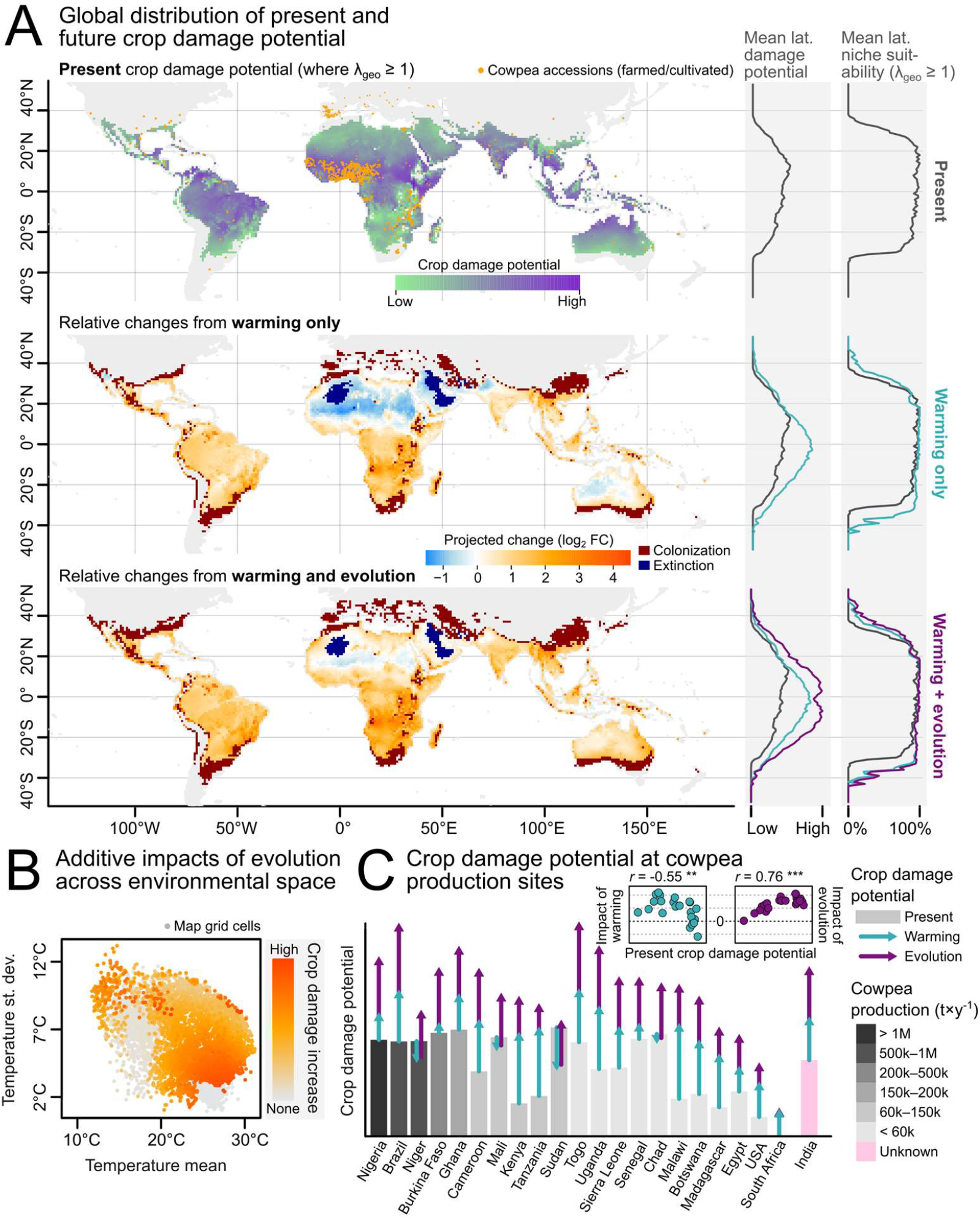
The contribution of evolution in forecasts of global crop losses under climate warming. (A) The top map displays the present-day crop damage potential as a baseline using ancestral (29 °C evolution regime) trait values, with overlaid points representing cowpea (*Vigna unguiculata*) accessions sourced from cultivated habitats. The lower maps show the log_2_fold-change in crop damage relative to the present-day baseline under future climate warming (SSP3-7.0, years 2081–2100), from direct, plastic effects of warming alone (middle) and from warming combined with thermal adaptation (bottom). Dark red and dark blue regions indicate novel colonizations (*λ*_geo_ goes ≥ 1) and localized extinctions (*λ*_*geo*_falls < 1). Marginal plots show crop damage potential and occupancy probability (*λ*_geo_≥ 1) averaged across latitude for the present-day baseline (grey), warming-only scenario (teal) and warming + evolution scenario (purple). (B) The additive impact of evolution on future crop damage mapped across a two-dimensional environmental space defined by recent (2018–2024) temperature means and standard deviations. (C) Projected crop damage for the world’s top cowpea-producing nations (comparable production data lacking for India) at reported cowpea accession sites, weighted by accession frequency within each country. Bars indicate present-day baseline damage potential, and arrows show the projected change in damage under warming alone (teal) and with added effects of pest evolution (purple). Inset scatterplots in (C) show the correlation between present-day damage and the changes driven by warming or evolution, with values corresponding to the length of the arrows in the bar chart. All projections are performed at a 1° spatial resolution, with spatially and seasonally resolved climate warming patterns.

**Table 1.** Projected crop damage potential under present and future climates (SSP3-7.0). Projected crop damage potential with and without local evolutionary adaptation, summarized as (A) damage potential aggregated over grid cells, (B) the relative impacts of gradual rate changes versus extinction risks/colonization opportunities based on thermal niche viability, and (C) the proportion of grid cells where evolution flips a warming-driven decrease in damage into an increase above present levels. Present = ancestral populations under current climates. Warming = ancestral populations under future climates. Warming + evolution = best-performing populations (highest *λ*_*geo*_) per grid cell under future climates (SSP3-7.0). Grid cells with *λ*_*geo*_ < 1 contribute zero damage potential in their scenario. “Cowpea producers” signify grid cells with cowpea accessions from cultivated/farmed areas in leading cowpea-producing countries (those listed in Boukar et al. 2019, plus India). The contribution of each cowpea producing grid cell was weighted by the number of unique accession coordinates in that specific cell. Percentages are based on averages over a common mask (all grid cells within a certain category).

| **A** |  | Crop damage rate (in growth rate; g/h/mother) |  |  | Change relative to present (%) |  | Evolution / warming |
| --- | --- | --- | --- | --- | --- | --- | --- |
| Geographic scope | Aggregation | Present | Warming | Warming + evolution | Warming | Warming + evolution | Fold ratio |
| Global | Unweighted grid cell mean | $2.88 \times 10^{-4}$ | $4.55 \times 10^{-4}$ | $6.18 \times 10^{-4}$ | +58.2% | +114.6% | 1.97x |
| Global | Area-weighted grid cell mean | $3.36 \times 10^{-4}$ | $5.31 \times 10^{-4}$ | $7.18 \times 10^{-4}$ | +57.8% | +113.4% | 1.96x |
| Cowpea producers | Accession-weighted mean | $8.00 \times 10^{-4}$ | $1.03 \times 10^{-3}$ | $1.49 \times 10^{-3}$ | +29.2% | +86.8% | 2.98x |

| **B Attribution of change in global mean damage rate** |  |  |  |  |  |  |
| --- | --- | --- | --- | --- | --- | --- |
| Scenario | Component | n grid cells | % of all grid cells | Absolute change | % of present | % of total change |
| Warming only | Gradual changes ($\lambda$ remains $\geq 1$) | 5,108 | 44.0% | $1.37 \times 10^{-4}$ | +47.7% | +82.0% |
| Warming only | Decrease from local extinction ($\lambda$ falls $< 1$) | 206 | 1.8% | $-7.51 \times 10^{-6}$ | -2.6% | -4.5% |
| Warming only | Increase from local colonizations ($\lambda$ goes $\geq 1$) | 1,037 | 8.9% | $3.77 \times 10^{-5}$ | +13.1% | +22.5% |
| Warming + evolution | Gradual changes ($\lambda$ remains $\geq 1$) | 5,143 | 44.3% | $2.83 \times 10^{-4}$ | +98.2% | +85.6% |
| Warming + evolution | Decrease from local extinction ($\lambda$ falls $< 1$) | 171 | 1.5% | $-6.02 \times 10^{-6}$ | -2.1% | -1.8% |
| Warming + evolution | Increase from local colonizations ($\lambda$ goes $\geq 1$) | 1,340 | 11.5% | $5.34 \times 10^{-5}$ | +18.5% | +16.2% |

| **C Scenarios where evolution flips a warming-driven crop damage decline to an increase** |  |  |
| --- | --- | --- |
| Subset | n grid cells | % of common mask |
| Grid cells where warming alone decreases damage (ancestor → ancestor) | 1,533 | 13.2% |
| ... of which evolution lifts damage above present (flip) | 997 | 8.6% |
| Flip rate (of grid cells where warming alone reduces damage, % with evolution-driven damage above present): 65% |  |  |
| Of these 997 flipped grid cells, 49 (4.9%) are in two top-producers of cowpea, Nigeria + Niger — 3.2× their 1.5% share of global modelled land. |  |  |

Adding evolution to the projections strongly amplified crop loss forecasts (Fig 5A), shifting the area-weighted increase in damage potential to +113%, a near doubling from the projected average impact of warming alone (Tab 1A). Moreover, warming is—in the absence of evolution—projected to reduce crop damage potential in 1,533 grid cells across the globe (Tab 1C). In 65% of these, adding evolution to the projections reverses the effect (Tab 1C), turning a projected decrease into an increase (blue to red in Fig 5A). These reversals are concentrated in the largest cowpea-producing regions: per unit of land area, they occur 3.2 times as often in Nigeria and Niger as the global average (Tab 1C).

Indeed, the impacts of evolution under future climate change are generally predicted to be highest in warm regions where *C. maculatus* already has high present-day crop damage potential, and where cowpea production is high (Fig 5). Accession-weighted projections showed that while the relative pest damage impact of warming alone is low at an average of +29% across cowpea production sites, incorporating pest evolution triples the total impact, with future mean crop damage potential increasing +87% above current baselines. Notably, a negative correlation (*r* = −0.55, P < 0.01, Fig 5C) between present-day crop damage risk and expected future changes driven by warming among top cowpea-producing countries suggests that warming might alleviate crop damage in current high-risk regions in the absence of evolution. But this might give a false sense of security; there is also a strong positive correlation (*r* = 0.76, P < 0.001, Fig 5C) between a country’s present-day risk and the projected warming-driven increase in crop damage owing to evolution, identifying pest evolution as a hidden potential threat to future crop yields.

## Discussion

Our findings show that pest evolution can amplify the threat climate change poses to global agriculture via several distinct routes. First, and most intuitively, pest adaptation to heat can increase intrinsic population growth under warming (Figs 2–5), including in regions where warming would otherwise reduce or eliminate the demographic growth potential of ancestral genotypes. Second, adaptation to cold expands the latitudinal range-limits of the thermal niche (Figs 4,5), potentially allowing pest insects to more efficiently track host plant cultivation zones that are rapidly shifting poleward. Third, evolution of specific life-history strategies favored under climate change can increase crop consumption rates beyond what would be predicted from pest fitness gains alone. In the hot-adapted *C. maculatus* populations, we found that fitness was increased mainly via increased fecundity, which itself was facilitated by evolved increases in adult body mass. Since larvae must feed more to pupate at a larger size, and since host consumption scales with biomass production rather than with offspring number, crop damage potential under climate warming was amplified by the hot-adapted beetles’ life-history strategy (Fig 3).

The observed evolutionary dynamics challenge the assumption that demographic declines due to warming would alleviate pest impacts in already-warm regions^2,6,9,55–57^. Instead, our projections show that thermal adaptation might not only counteract warming-induced declines in fitness but sometimes even rescue populations from temperature-driven extinction. Indeed, incorporating evolution in the projections frequently flipped damage reductions into substantial increases, particularly in major cowpea-producing regions (Fig 5, Tab 1). Thus, in areas like sub-Saharan West Africa, where cowpea production constitutes a vital source of both food security and agricultural income^58,59^, the ability of *C. maculatus* to adapt and maintain high crop consumption rates—even as climate warming pushes temperature distributions toward new extremes— constitutes an alarming potential amplifier of crop losses, adding to the socioeconomic costs of extreme climates themselves^6,31,44^.

Notably, even highly seasonal thermal environments with relatively low yearly mean temperatures can temporarily become extreme as warming raises summer temperatures^60^. However, as seen in our results, the negative impact of such extreme summer events can be partly offset by a widening of the pest’s growth season (Fig 2D)^60^. Thus, growth season expansion can be a major driver of increases in annualized crop-damage, as it prolongs the window where warm conditions sustain high host consumption rates (Fig 2C). Additionally, for species like *C. maculatus* that do not exhibit winter diapause, seasonal environments with prolonged cold periods frequently fall outside the habitable thermal niche (*λ*_*geo*_ < 1), but warming shortens these periods, potentially enabling poleward spread into new crop producing regions (Figs 2D,5A).

This influence of a shifting growth season on crop damage potential underscores the importance of a temporally dynamic modelling framework. Approaches that fix the pest’s active season (e.g., confining it to the crop’s current growing period^6^, even though that may itself shift under warming^61^) cannot fully capture how seasonality and climate change interact to determine crop damage. The seasonal dynamics in our simulations also highlight how large relative differences in fitness do not necessarily translate into substantial differences in crop damage; evolved fitness increases in cold environments contribute little to total absolute crop damage potential because biological rates (and thus host consumption) remain low at low temperatures, so the yearly impact is dominated by what happens during the warm season (Fig 2B,C). Instead, cold adaptation raises global crop damage potential mainly by pushing *λ*_*geo*_ above 1 in high-latitude environments where, paradoxically, damage potential mainly accrues during the warm part of the year (Figs 2,4).

An interesting implication of our findings is that accurately forecasting pest responses to climate change requires considering not only *if* pests will adapt to new temperature distributions, but also *how* the specific life-history pathways that underlie fitness outcomes shape agricultural impacts^12^. The evolution of thermal sensitivity (i.e., shifts in the *shape* of reaction norms) may be relatively constrained^62–64^ but importantly, our results demonstrate that such evolution is not a requisite for thermal adaptation, which here was achieved mainly through changes in allocation patterns between different life-history components (Figs 1–3). In the case of *C. maculatus* evolving larger body sizes under warm temperatures, this matters greatly for forecasting. While larger insect body sizes with warming are not the universal norm^65–67^ (but common in the pest-rich insect orders of Coleoptera, Lepidoptera, and Orthoptera^68^), thermodynamic effects on metabolic rate^69^ can increase overall energy flux and provide a higher capacity for biomass production, potentially allowing larger body sizes (generally linked to higher fitness^67^) to evolve under warm conditions^12,70–73^. Alternatively, warm temperatures might select for increased feeding rates^74^ just to keep up with heightened metabolic demands^19^ while body sizes remain constant or even decrease as a result of selection^75^—particularly under limited resource availability^22^. In both scenarios, evolved increases in pest consumption rates constitute an additional agricultural threat layered on top of the direct metabolic effects of warming^6^. Thus, amplified crop damage potential as a consequence of adaptive responses to warming may prove a general phenomenon in ectothermic crop pests, particularly under conditions of abundant food availability typical of cultivated landscapes^21–23^.

Given the computational scale of our projections, our modelling framework naturally involves certain methodological choices for tractability. Still, it provides an exceptionally high eco-physiological resolution and avoids the pitfalls of treating fitness as a single, instantaneous function of temperature by decomposing responses into separate life-history processes^28^. The approach nonetheless makes some simplifying assumptions—primarily that temperature responses aggregate additively (multiplicatively for viability) over time and that each can be described as a continuous function of temperature throughout ontogeny^39,76–82^. It should also be noted that, while local weather conditions can provide strong explanatory power for predicting temporal variation in insect biomass^83^, temperature is not the only driver of pest responses^9^, and its impacts can be further modulated by behavioral processes like thermoregulation^11,84^. Thermoregulation may relax selection pressures from rising temperatures and impede physiological adaptation to heat^85^, but it can instead amplify crop losses by buffering the pest’s exposure to harmful temperatures^11^. Yet, *C. maculatus* is most destructive in the homogenous environments of dried bean storages^43^, where opportunities for thermoregulation might be limited. By omitting density dependence, we also avoid assuming carrying capacities and other regulatory parameter values; our projected outcomes therefore represent physiological *crop damage potential* and *thermal niche suitability* rather than absolute rates. Importantly, a single cowpea can support the development and successful adult emergence of over ten *C. maculatus* beetles, but the emergence of just a single beetle makes the bean unsuitable for human consumption. Thus, our unlimited growth metrics are arguably most relevant for predicting agricultural impacts; by the time pest density becomes limiting to population growth, the damage is already done. Finally, it should be noted that we map evolved responses onto projected climates rather than simulating evolution dynamically. As such, our projections are best interpreted as showing the capacity for evolution to shape future pest responses rather than absolute outcomes, which in reality will be influenced by a wide array of factors including gene flow^86^, trade-ofs^87^, the pace of warming relative to adaptation^88,89^, and temporal variation in selection pressures^90,91^. On the other hand, our three evolution regimes undersample the possibility-space of evolutionary responses, and adaptation to constant temperatures may be suboptimal under variable settings—from which perspective our estimates are conservative with regard to the beetles’ evolutionary potential.

Together, our results highlight that accurately forecasting pest responses to climate change requires considering not only changes in pest fitness, but also the specific life-history adaptations underlying them. In our *C. maculatus* projections, evolution rescued populations from suffering reduced fitness under warming while also amplifying their destructive potential through a life-history strategy involving increased larval growth and host consumption rates—plausibly commonplace pest adaptations to a warming world. Notably, the evolutionary amplification of crop damage increases was particularly pronounced in the major crop-producing regions of sub-Saharan Africa. We conclude that excluding the possibility of adaptive evolution in forecasts of future pest impacts may underestimate how climate change threatens food production, particularly in warm regions with already limited food security.

## Methods

### Experimental evolution

Experimental evolution lines of the seed beetle *Callosobruchus maculatus* were established in 2013 from three geographically distinct ancestral populations (Brazil, Yemen, and California, USA) that had been maintained under common-garden conditions at 29 °C since 2002 at Uppsala University, Sweden. Each ancestral population was split into two novel thermal environments— 23 °C and 35 °C—with two replicate lines per geographic origin per treatment, yielding six lines in each evolution regime. The founding ancestral lines were maintained under 29 °C as they had been since before the establishment of the new evolution regimes. The experimental evolution lines maintained at 23 °C and 35 °C had undergone approximately 90 and 130 generations of selection, respectively, by the time the assays were performed (the discrepancy reflects the effect of temperature on generation time). Throughout the experiment, populations were reared at 55% relative humidity on dried *Vigna unguiculata* (cowpea) beans. Detailed rearing protocols are described previously^12,92^.

### Thermal performance assays

Thermal performance was assayed across five constant temperatures (17, 23, 29, 35, and 37 °C) at 55% relative humidity. After two generations of common garden at 29 °C, virgin mating pairs were isolated in individual Petri dishes provisioned with *V. unguiculata* beans and transferred to the designated temperature treatment. Three response variables were recorded: lifetime reproductive success (LRS; total offspring count), development time (days from first oviposition to first adult offspring emergence), and offspring dry weight. Measurements were replicated across all combinations of assay temperature, evolution regime, geographic origin, and replicate line, with a minimum sample size of n = 10 at the lowest level of replication (n^tot^ = 1,211 parental pairs). Paired individuals that proved to be of the same sex upon post-experiment dissection were excluded from analyses.

### Bayesian fitting of thermal performance curves

Thermal performance curves (TPCs) were fitted as Bayesian non-linear models in Stan^93^ via the R^94^ package brms^95^. Three separate models were specified for fecundity and viability, development rate, and growth rate.

#### Fecundity and viability

Lifetime reproductive success was modelled as a two-component process comprising (1) viability—the probability that a pair produces any offspring at all—and (2) fecundity conditional on viability.

Viability was modelled on the logit scale as a quadratic function of temperature:

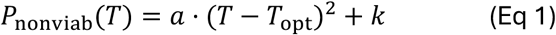

where *a* is the curvature parameter governing the temperature dependence of becoming non-viable (constrained to be positive; the probability increases at extreme temperatures), *T*_opt_ is the temperature at which viability is maximized (i.e., non-viability is minimized), *k* is the minimal log-odds of non-viability at *T*_opt_, and *T* is the ambient temperature.

Fecundity conditional on viability was modelled as a generalized bell-shaped function of temperature:

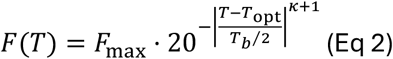

where *F*_max_is the maximal offspring number, *T*_opt_is the temperature that maximizes fecundity, *T*_b_is the temperature breadth over which fecundity exceeds 5% of its maximum (as set by the base 20 since 20^-1^ = 5%), *κ* is a kurtosis parameter controlling the “peakedness” of the curve, and *T* is the ambient temperature.

All parameters in Eq 2 were estimated separately for each geographic origin regime combination, with the exception of the kurtosis parameter *κ*, which was estimated jointly across cold- and hot-adapted lines but separately for the ancestral regime to aid model convergence. Random estimates for each replicate line were included for *T*_opt_, *F*_max_, and *T*_b_, whereof the random effect on *F*_max_ was implemented on the log-scale. Viability parameters (Eq 1) were estimated per evolution regime without origin-specific effects or random intercepts to aid model convergence. The response was modelled with a zero-inflated negative binomial error distribution, where the zero-inflation component captures non-viability (Eq 1) and the count component captures fecundity (Eq 2), with temperature- and regime-specific dispersion parameters due to the data’s non-homoscedasticity on the log scale.

#### Development

Development rate (inverse of development time in days) was modelled using the Lobry–Rosso– Flandrois (LRF) function:

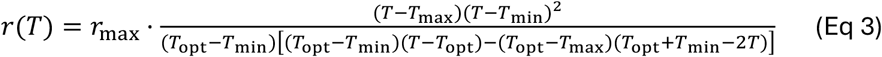

with *r*(*T*) = 0 for *T* < *T*_min_ or *T* > *T*_max_. Here, *r*_max_ is the maximal development rate, and *T*_min_, *T*_opt_, and *T*_max_ are temperature parameters (minimum, optimum, and maximum, respectively). The error distribution was assumed to be lognormal. Parameters were estimated for each geographic origin × evolution regime combination, with random intercepts per replicate line for each temperature parameter and *r*_max_. The random effect on *r*_max_ was implemented on the log-scale.

#### Growth

Additive growth rate (offspring dry weight divided by development time) was modelled with the same LRF function (Eq 3) and random effects structure used for development rate. To improve identifiability, Gaussian priors (SD = 1) on *T*_min_ and *T*_max_ were centred on the posterior modes from the corresponding development rate model for each geographic origin × evolution regime combination.

#### Global models

In addition to the origin-specific models described above, analogous “global” models were fitted for each trait (development rate, growth rate, and fecundity/viability) with the evolution regime-specific temperature dependence averaged across geographic origins. Instead, random intercepts were fitted at the replicate and replicate × temperature treatment level (replicate lines are nested within origins, such that the replicate-level random intercepts absorb origin-level variation). These global models were used for the global map projections.

#### Priors and assessment of fit

When possible, we specified informative priors for the nonlinear model parameters (Table S1). For instance, the curvature parameter *a* was bounded to be positive, and temperature parameters were constrained via Gaussian distributions centered on biologically plausible values (e.g., assuming that *T*_opt_for development has a ∼95% probability to fall between 27 °C and 43 °C). Posterior predictive checks confirmed that the generative models reliably approximated the observed experimental data distributions (Supporting Information; Figs S8–S13).

### From thermal responses to hourly rate functions

To become combinable with hourly temperature data, all trait-specific responses were converted to hourly rate functions.

#### Development and growth rate

Daily rates obtained from the fitted TPC models (Eq 3) were divided by 24 to yield hourly rates:

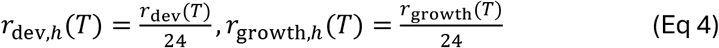

#### Oviposition rate

A temperature-dependent conversion factor relating short-term oviposition rate to lifetime reproductive success was derived empirically from published data on 18-hour offspring production and LRS across temperatures^12^. The log-ratio of short-term to lifetime output was approximately linear with temperature, and a batch correction was applied to unify the ancestral and evolved lines using a population scored in both experimental batches (Figs S14,S15; see also Burc et al. 2025^12^). The conversion was calculated separately for ancestral and selected (cold- and hot-adapted) lines. The hourly oviposition rate is:

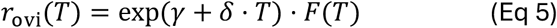

where *γ* and *δ* are the intercept and slope of the log-linear temperature relationship (estimated separately for ancestral and selected lines), and *F*(*T*) is the fecundity function (Eq 2).

#### Viability rate

The probability of non-viability (Eq 1) was converted to an hourly viability rate by raising the viability probability to the power of the expected hourly development rate, distributing the probability evenly over the expected development period:

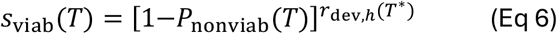

where *T*^*^ = max(17, min(37, *T*)), which clamps the boundaries of development rate to its values at the extremes of the assay range (17 and 37 °C) to prevent nonsensical extrapolation (a development rate of zero would erroneously predict perfect fertility since *x*^0^ = 1).

#### Fecundity-mass scaling

To account for the effect of parental body mass on reproductive output, a log-linear relationship between dry mass (in grams) and relative fecundity was estimated from the experimental data at 29 °C (the benign ancestral rearing temperature), with evolution regime as a covariate (Fig S3):

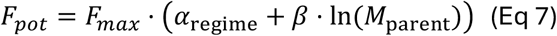

where *F*_max_ is the maximum fecundity from the fitted curve (Eq 2), α_reg me_ is a regime-specific intercept, β is a mass-scaling exponent, and *M*_parent_ is the body mass of the ovipositing parent. Offspring counts were normalized within origin and regime (at 29 °C) before fitting, so that the regression isolates the multiplicative effect of individual mass on fecundity relative to the expected value captured by Eq 2.

### Life-cycle simulation algorithm

We constructed an algorithm for projecting individual-level fitness and crop damage rates over hourly temperature time-series from the hourly rate functions (Eqs 4–7). For each day of a given focal year, the simulation assumes the emergence of an adult beetle (“focal female”) and projects its reproductive output forward in time. Each daily batch of eggs deposited by the focal female is tracked as an independent offspring cohort so that offspring laid on different days experience distinct temperatures. The algorithm proceeds as outlined below.

#### Pre-computation

All four rate functions (development, growth, viability, oviposition) were evaluated for every hour of a multi-year temperature series spanning the focal year and surrounding years for backward (parental mass) and forward (offspring development) integration. Series of cumulative sums were constructed for each process, enabling outcomes over any time window to be efficiently computed by subtraction. For viability, the cumulative sum was computed on the log scale since it is a multiplicative process. Offspring cohort computations extended beyond the focal year to accommodate offspring produced late in the season whose development continues into the following year.

#### Step 1: Parental mass and fecundity scaling

The body mass of the focal female was estimated by backward-in-time integration of the growth rate function over one developmental period preceding emergence. Starting from the hour of emergence, the cumulative development rate was tracked backward in time until it reached 1.0 (i.e., completed development), and the cumulative growth rate over that window gives the parental mass:

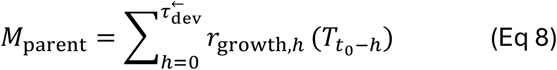

where t_O_ is the hour of adult emergence and 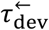 is the backward-calculated development time, defined analogously to the forward development time calculation (Eq 10). The maximum potential fecundity of the focal female was then adjusted according to its mass via Eq 7.

#### Step 2: Oviposition

Beginning at adult emergence, all females were assumed to be fertilized, in line with the rapid female mating rates, lack of sperm limitation, and high male re-mating rates observed in *C. maculatus*^96–98^. Hourly egg laying was modelled using the temperature-dependent oviposition rate (Eq 5), modulated by an extrinsic mortality rate to get the expected realized oviposition output (Eq 13):

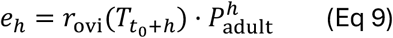

where *e*_*h*_ is the expected number of eggs deposited at hour *h* of the focal female’s adult life and 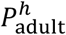 is the cumulative survival probability of the adult at hour *h* based on a pre-defined extrinsic mortality rate (defined below). Oviposition continued until either the fecundity pool was exhausted or adult survival dropped below 1%.

#### Step 3: Offspring development time

For each offspring cohort (corresponding to eggs laid on a specific day of the focal female’s reproductive period), the cumulative hourly development rate was tracked forward from the hour of oviposition until it reached 1.0, indicating completion of juvenile development after *τ*_dev_ hours:

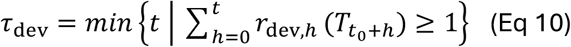

#### Step 4: Offspring mass

The cumulative hourly growth rate integrated over the cohort-specific development period (Eq 10) gives the expected dry weight of emerging offspring:

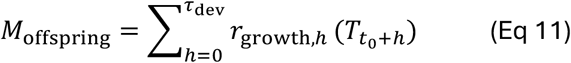

#### Step 5: Offspring viability

The product of hourly viability rates (Eq 6) over the cohort-specific development period yields the probability of offspring remaining viable at adult emergence:

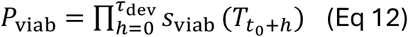

#### Step 6: Extrinsic mortality

Because the extrinsic (i.e., not temperature driven) mortality is unknown for natural populations, three alternative scenarios were considered, corresponding to uniform mortality rates generating a mean adult lifespan of 5, 10, and 20 days. Juveniles reside within individual seeds and are thus less exposed than adults, and we therefore set the expected juvenile lifespan to twice the adult lifespan in each scenario. Hourly survival probabilities were derived from these lifespans assuming exponential decay:

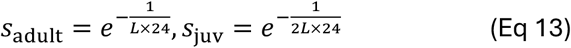

where *L* is the expected mean adult lifespan in days. The adult survival rate modulates the expected oviposition output as 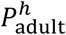 in Eq 9. The juvenile survival rate gives the probability of an individual from an offspring cohort remaining viable after the cohort’s specific development period:

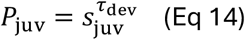

#### Step 7: Per-cohort fitness components

For a focal female emerging on day *i* of the temperature time-series, steps 1–6 yield a sequence of daily offspring cohorts indexed by the day *d* of the focal female’s reproductive period (on which their eggs were laid). Each cohort *d* gets assigned an expected egg count *e*_d_ (based on cumulative temperature-driven oviposition rates given in Eq 9 and accounting for adult extrinsic mortality), development time *τ*_dev,*d*_ (Eq 10), dry mass at emergence *M*_offsprng,*d*_ (Eq 11), thermal viability *P*_viab,*d*_ (Eq 12), and juvenile extrinsic survival *P*_juv,*d*_(Eq 14). The total generation time (*G*) of cohort *d* is calculated as the elapsed time from focal female emergence to offspring adult emergence:

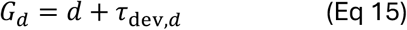

The expected number of viable adult offspring contributed to the population by cohort *d* is:

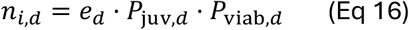

Thus, Eq 16 combines the per-cohort egg count, juvenile survival, and thermal viability into a single metric.

#### Step 8: Linear offspring production and crop damage rates

From Eqs 15 and 16, we calculate both a linear rate of per-capita offspring production (monotonically related to fitness; Eq 17) and crop damage (Eq 18) by weighting the per-cohort contributions by the inverse of their generation time. These metrics represent the time-averaged fluxes of surviving female offspring (*J*_offspring,*i*_) and crop biomass consumption (*J*_damage,*i*_) resulting from a focal female emerged on day *i*.

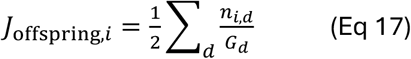

In Eq 17, the factor of *½* converts total surviving offspring to surviving females. Thus, *J*_offspring,*i*_ represents a female’s net reproductive rate divided by the offspring-weighted harmonic mean of her offspring cohorts’ generation times (each cohort is divided by its own generation time). The crop damage rate is an analogous expression (but which includes male offspring as well) weighted by each offspring’s expected biomass (which predicts host consumption—beetle mass produced is approximately 3.5x the host mass consumed^12^; R^2^ = 0.93, Fig S7):

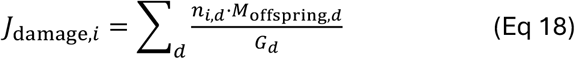

#### Step 9: Demographic rate of increase

We also calculated long-term geometric population growth as the finite rate of increase (λ). The net reproductive rate (*R*_0_) of a focal female emerging on day *i* is:

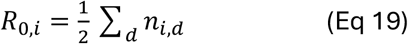

and the offspring-weighted mean generation times of those female offspring was calculated as:

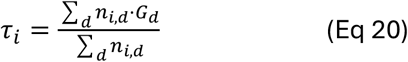

We then calculated the daily finite rate of increase using an easily computable approximation of the Euler-Lotka equation, which assumes negligible variance in generation times among individuals within the same cohort (justified by the brief reproductive period in comparison to the developmental period):

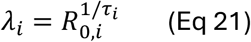

### California case study

We retrieved hourly temperature records (2018–2023) from 20 California weather stations (CIMIS^99^) spanning a broad thermal gradient (8.3° latitude, >10 °C in mean temperature), chosen because California is both the origin of one of our genetic backgrounds and a major US cowpea producer^51^. Using a custom algorithm that preserves realistic daily and seasonal cycles and residual autocorrelation (*SI Extended Methods*), we simulated hourly regimes at each site and projected them to 2100 under three warming scenarios (0.02, 0.04, and 0.06 °C yr^-1^).

For each geographic origin and site, we ran our year-long simulation algorithm under present-day conditions and year 2100 warming scenarios. We first evaluated the full, integrated phenotypes of the ancestral, cold-adapted, and hot-adapted regimes. To then quantify the isolated contribution of individual life-history traits to the overall resulting impact on population fitness and crop damage, we constructed artificial phenotypes *in silico*. For these, a single focal trait (development, growth, viability, or fecundity) was parameterized according to an evolved regime’s (hot- or cold-adapted) thermal reaction norm, while all other life-history traits were constrained to the ancestral response.

To evaluate seasonal and site-specific dynamics in climate warming responses, we extracted per-capita offspring production rates (monotonically related to fitness; Eq 17) and linear biomass consumption rates (representing crop damage potential; Eq 18) at each day of the year across the different scenarios. The resulting 1-day resolution time-series allowed us to calculate temporally resolved differences in responses among evolution lines and geographic locations. We also calculated site-specific relative differences in yearly average rates relative to present-day baselines, allowing us to determine the additional impacts of evolution beyond strictly plastic responses to temperature change and partition them across life-history traits. Finally, to test the sensitivity of our projections to demographic and environmental assumptions, we repeated the analyses under different constant rates of extrinsic mortality (corresponding to expected adult lifespans of 5, 10, and 20 days, with twice as long juvenile survival expectancy) and using daily mean temperatures in place of hourly thermal fluctuations.

### Global analysis

#### Projections

To extend our projections to a global scale, we applied our life-cycle simulation algorithm to each cell of a 1° global grid using the “global” (evolution regime-only) thermal responses and the intermediate extrinsic mortality. Between latitudes 60 °S to 60 °N, each grid cell was assigned an hourly temperature time-series for a present-day reference period (2018–2024) and for an intermediate-to-warm future climate scenario. The hourly air temperature data were obtained from the ERA5-Land reanalysis^52^ with a resolution of 1° using the R package *ecmwfr*^100^, which provides access to the Copernicus Climate Data Store API. These served as the baseline for estimating present-day temperature responses in fitness and crop damage rates. We then generated and simulated responses under future temperature regimes using a mean-shift approach, whereby grid cell-specific projected changes in monthly mean temperature from global climate models were applied to the historical ERA5 hourly time-series. These spatially resolved climate change projections were obtained from WorldClim v2.1^53^ (SSP3-7.0 scenario for the period 2081-2100; ACCESS-CM2 global climate model).

#### Plastic and evolutionary responses to climate warming

To determine the spatial distribution of both direct impacts of climate warming and the added impacts of evolutionary responses, we evaluated three primary scenarios for every grid cell: (1) a present-day baseline using the ancestral thermal reaction norms; (2) a “warming only” scenario calculated from ancestral plastic trait responses under future climates; and (3) a “warming and evolution” scenario. For the evolutionary scenario, we identified the “winning” thermal strategy per grid cell, defined as the evolution regime yielding the highest geometric mean finite rate of increase (*λ*_*geo*_) under future climatic conditions. We then used the thermal reaction norms of this winning regime to calculate the crop damage potential within the focal grid cell. In all scenarios, grid cells where *λ*_*geo*_ fell below 1 were classified as non-viable as they cannot sustain a population indefinitely, and were therefore assigned a crop damage rate of zero.

For visualization, we mapped average present-day crop damage potential (in g day^-1^ mother^-1^, linearly correlated with crop consumption^12^: R^2^ = 0.93, Fig S7), as well as the log_2_ fold-change in crop damage for future scenarios (both with and without evolution) relative to this present baseline. We decomposed the global shifts in crop damage potential into three distinct demographic components: crop damage shifts within cells that remain viable both in present-day and future conditions, decreases in crop damage potential resulting from localized demographic collapses (“extinctions”, *λ*_*geo*_falls < 1) and increases in crop damage potential driven by new grid cells falling within the thermal niche of *C. maculatus* (“colonizations”, *λ*_*geo*_goes ≥ 1). Furthermore, we also identified “flip” zones: regions where climate warming alone would decrease crop damage potential due to decreases in demographic growth but where evolutionary adaptation reverses this trajectory, lifting projected rates above present-day levels. Finally, we quantified the additive effect of evolution (the absolute difference in damage potential between the “warming and evolution” and “warming only” scenarios) and mapped it across a two-dimensional environmental space characterized by present-day mean temperatures and temperature variability. This analysis was restricted to grid cells that remained viable in future scenarios with evolution.

#### Future impacts in crop-producing regions

To translate our physiological and demographic projections into agricultural risk, we overlaid the global crop damage rasters (where *λ*_*geo*_< 1 signifies zero crop damage potential) with cultivation data for cowpea (*V. unguiculata*), *C. maculatus*’ primary host crop. We queried the Genesys database^54^via the genesysr R-package^101^to retrieve global accessions, filtering for *V. unguiculata* entries with valid geographic coordinates. To restrict our analysis to environments of known agricultural relevance, we only retained samples collected from farm or cultivated habitats (MCPD collecting-source codes 20–28^102^). Accessions sharing exact spatial coordinates were collapsed to represent a single sampling location. We assigned each unique accession point a crop damage rate corresponding to the 1° grid cell wherein it was located, which we then aggregated to create an accession-weighted damage metric, where cowpea sampling-frequency acts as a proxy for crop production. We then averaged accession-weighted crop damage rates (present-day and future) across the top 40 cowpea-producing nations identified and ranked in Boukar et al.^44^ We also added India to this list, as it is a major cowpea producer but without comparable production data due to alternative reporting practices^103^. Ultimately, this resulted in 22 unique nations, after excluding countries with fewer than two accession sampling locations.

## Use of generative AI

Generative AI (Claude Opus 4.6-4.8, Anthropic; Gemini 3.1 Pro, Google) was used to refactor and optimize some of the code, and occasionally to explore alternative phrasings when drafting the manuscript. In all cases, the output was validated manually (e.g., against human-written code), and we hold full accountability for the manuscript’s intellectual content.

## Acknowledgements

We thank Milena Trabert for letting us use her *C. maculatus* illustrations. David Berger thanks the Swedish Research Council for Sustainable Development (Formas) for funding (grant no. 2022-01117). Loke von Schmalensee thanks the Swedish Research Council (Vetenskapsrådet) for funding (grant no. 2025-07932).

## Notes

### Competing Interest Statement

The authors have declared no competing interest.

