## Supplemental Information for "Pest evolution amplifies projected crop losses under climate change"

### Contents

Extended methods (p. 2)

Figures S1–S21 (pp. 3–23)

Tables S1–S5 (pp. 24–31)

References (p. 32)

### Extended methods: temperature time-series simulation

To investigate fine-scale pest dynamics under climate warming across diverse thermal environments, we leveraged the availability of high-resolution weather station data from California—the geographic origin of one of the three genetic backgrounds we used in the experimental evolution and a major US producer of cowpeas (*V. unguiculata*) (Johnson & Valero, 2003), *C. maculatus'* main host.

To simulate realistic, empirically grounded, high-resolution thermal regimes under different climate warming scenarios, we first retrieved hourly temperature data from 20 weather stations in California, over the years 2018–2023 via the California Irrigation Management Information System (CIMIS) (California Department of Water Resources, 2018). We chose a range of stations spanning 8.3° latitude with average temperature differences of more than 10 °C between warm and cold sites (Fig S16). In order to retain temporal structure at different timescales, we designed a custom temperature simulation algorithm, incorporating coarse patterns of daily and yearly temperature cycles and autocorrelations in the residual temperature variation to generate realistic hourly thermal regimes *in silico*.

First, we converted the hourly temperature data from °C to log K. For each of the 20 sites individually, we then decomposed the temperature time-series into three components: an average daily cycle, an average annual (seasonal) cycle, and residual variation. The average daily cycle for a site was calculated by first splitting each year into 73 batches of 5 days each and then taking the mean log-transformed temperature per batch for each hour of the day across all observations for that site, centering the values by subtracting their mean. This captures diurnal fluctuations independent of the absolute temperature and retains a seasonal structure in the diurnal cycles. The average annual cycle was estimated by fitting a periodic smoothing spline to the daily mean log-transformed temperatures (across all years) for that site. We then calculated residuals by subtracting the sum of the predicted daily and annual components from the observed temperature at each time point.

Next, we modelled the autocorrelation structure within each site's residuals using an ARIMA model with both a non-seasonal (hour-to-hour) and a seasonal (24-hour time lags) autoregressive component. We then simulated a new sequence of residuals from this fitted model. To preserve the original level of variability, these simulated residuals were rescaled to match the standard deviation of the site's actual residuals.

Last, we summed the three components and converted the result back to °C, generating simulated thermal regimes with realistic daily and yearly cycles, temperature averages and variance, and temporal structure (Figs S17–S21). To each of the 20 site's simulated hourly time-series we then applied three linear warming trends of 0.02, 0.04, and 0.06 °C $yr^{-1}$ over 75 years (present day to year 2100).

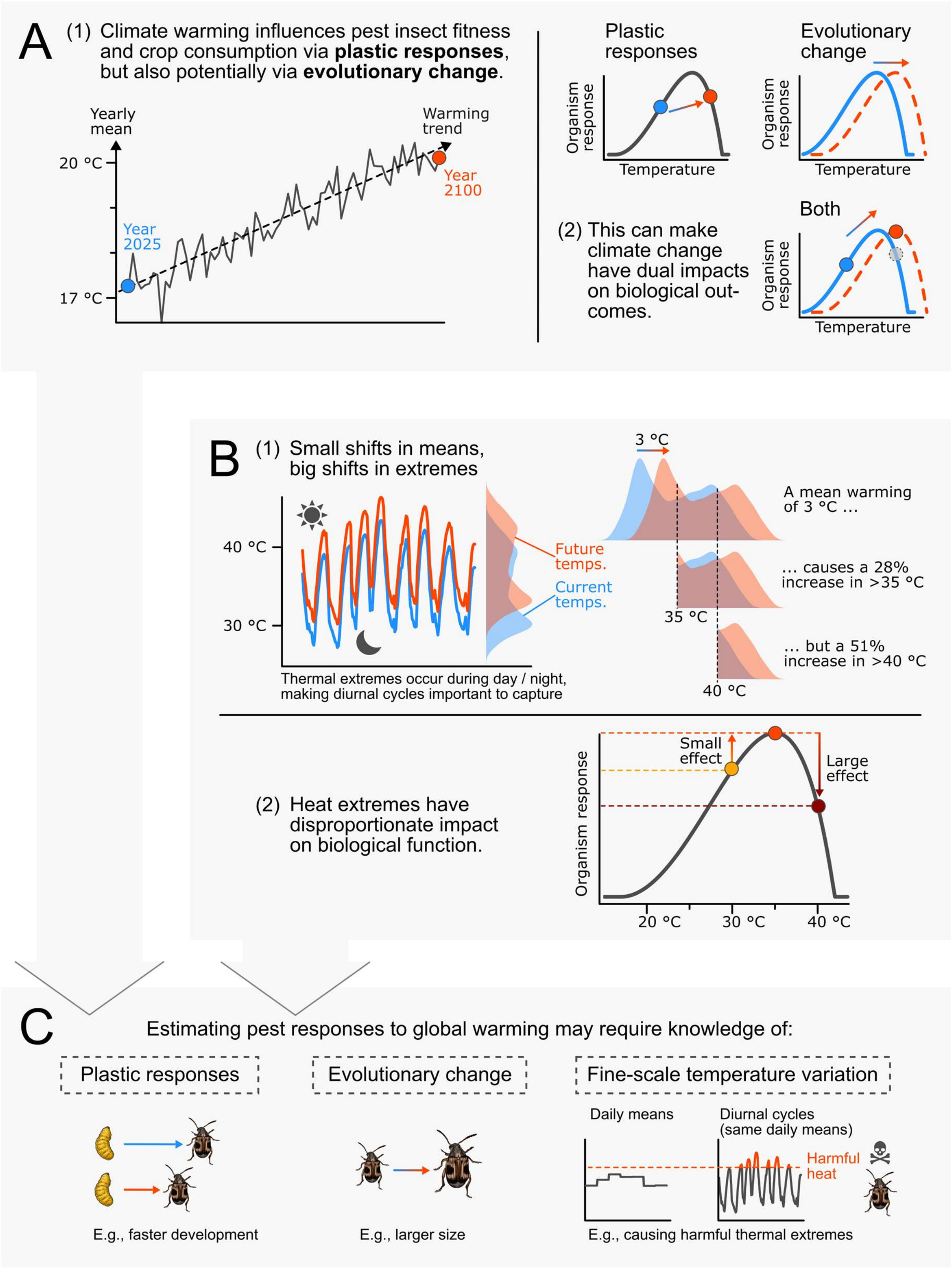

**Figure S1.** (A) Climate warming can influence pest insects through both immediate plastic responses and longer-term evolutionary changes. (B) Small increases in mean temperatures might lead to substantial increases in the frequency of thermal extremes. In turn, thermal extremes have disproportionate impacts on biological functions due to nonlinear reaction norms, underscoring the importance of fine-scaled temperature variation. (C) To accurately estimate pest responses to warming, it is therefore important to consider nonlinear plastic responses, evolutionary changes, and temperature variation at relevant scales, even when making long-term projections.

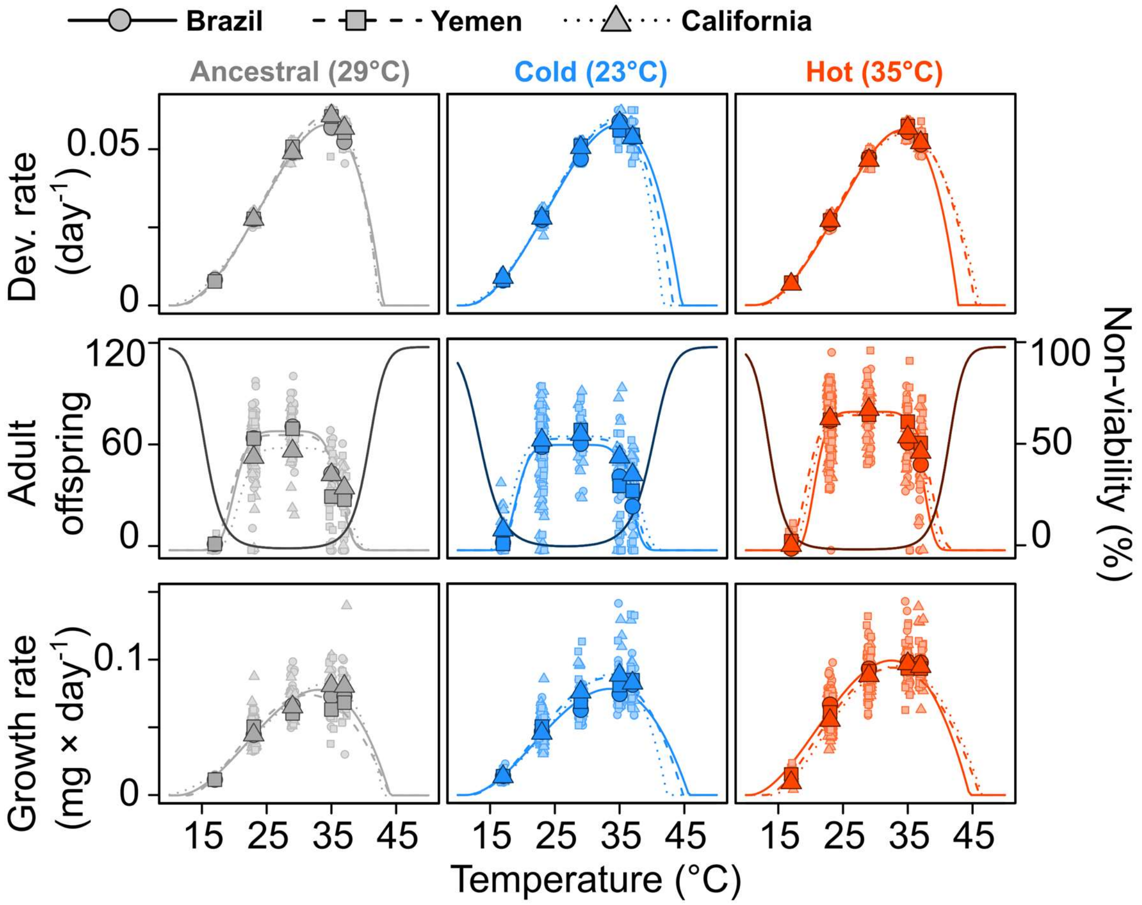

**Figure S2.** Raw data and fitted thermal reaction norms. Small points show observations; large points show temperature-specific averages for each line. For clarity, these averages exclude zeroes in the middle row (i.e., beetle pairs with no offspring), and the fitted excess probability (from a Poisson distribution) of having zero offspring is represented by the darker lines (right y-axis). The product of the probability of viability and the expected number of offspring (given viability) represents lifetime reproductive success (LRS).

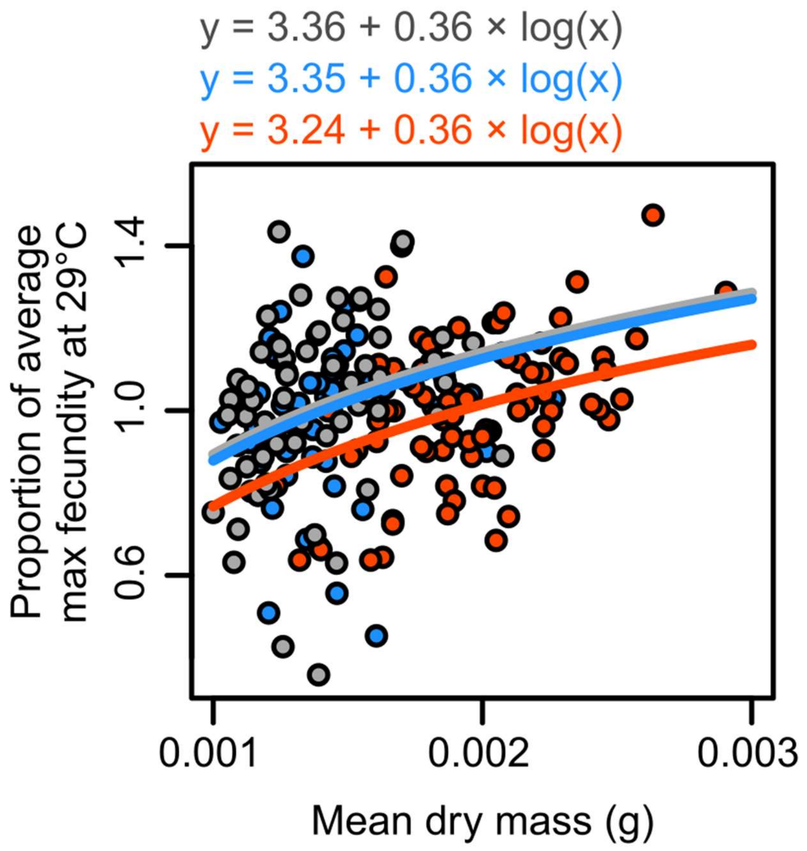

**Figure S3.** The effect of beetle size on relative maximal fecundity (scaled within evolution regime at 29°C). Grey represents ancestral lines, blue represents cold adapted lines, and red represents hot adapted lines. Text shows the corresponding fitted equations.

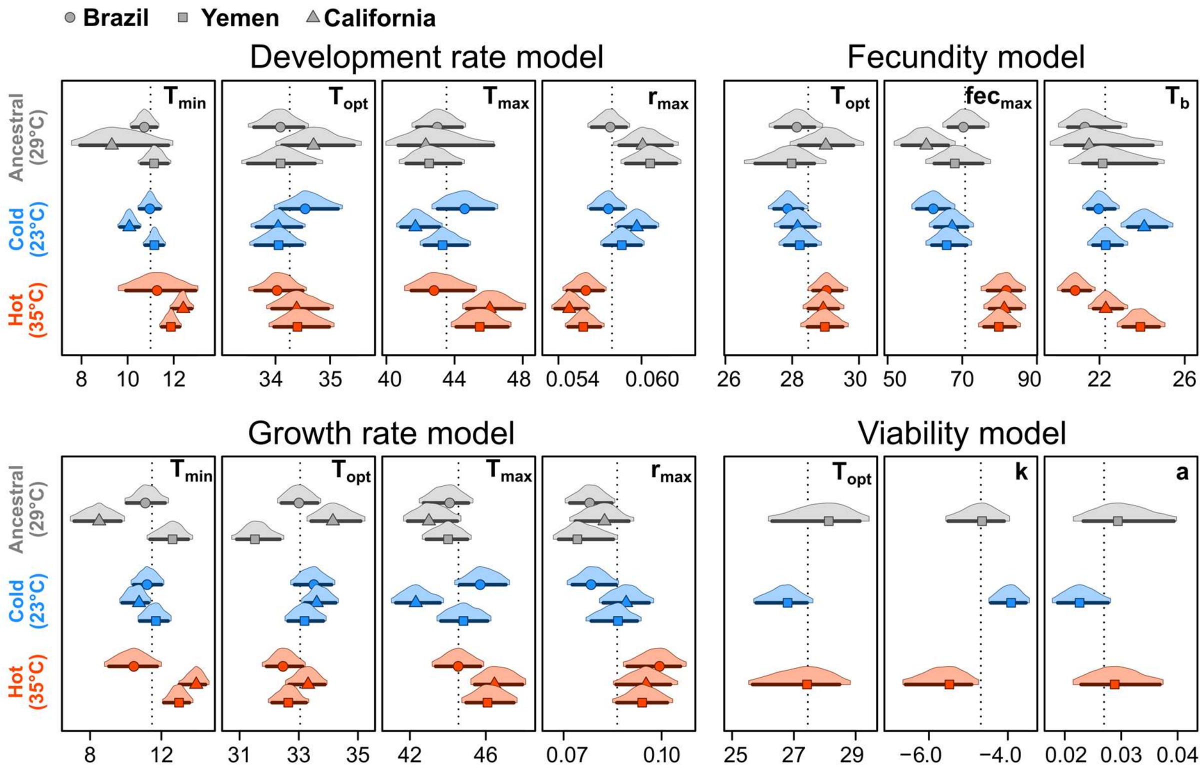

**Figure S4** Posterior distributions of thermal reaction norm parameters. Densities cover 95% highest posterior density intervals (HPDIs), lines show 85% HPDIs, and points represent posterior modes. $T_{min}$ = minimum temperature for a non-zero response; $T_{max}$ = maximum temperature for a non-zero response; $T_{opt}$ = the temperature where the response is maximized; $r_{max}$ and $fec_{max}$ = the maximal response; $T_b$ = difference between the maximum and minimum temperature where fecundity is > 5% of $fec_{max}$; $k$ = the log-odds of non-viability at $T_{opt}$; a = the curvature parameter governing the temperature dependence becoming non-viable, with higher values representing a more rapid decline in viability probability away from the optimum.

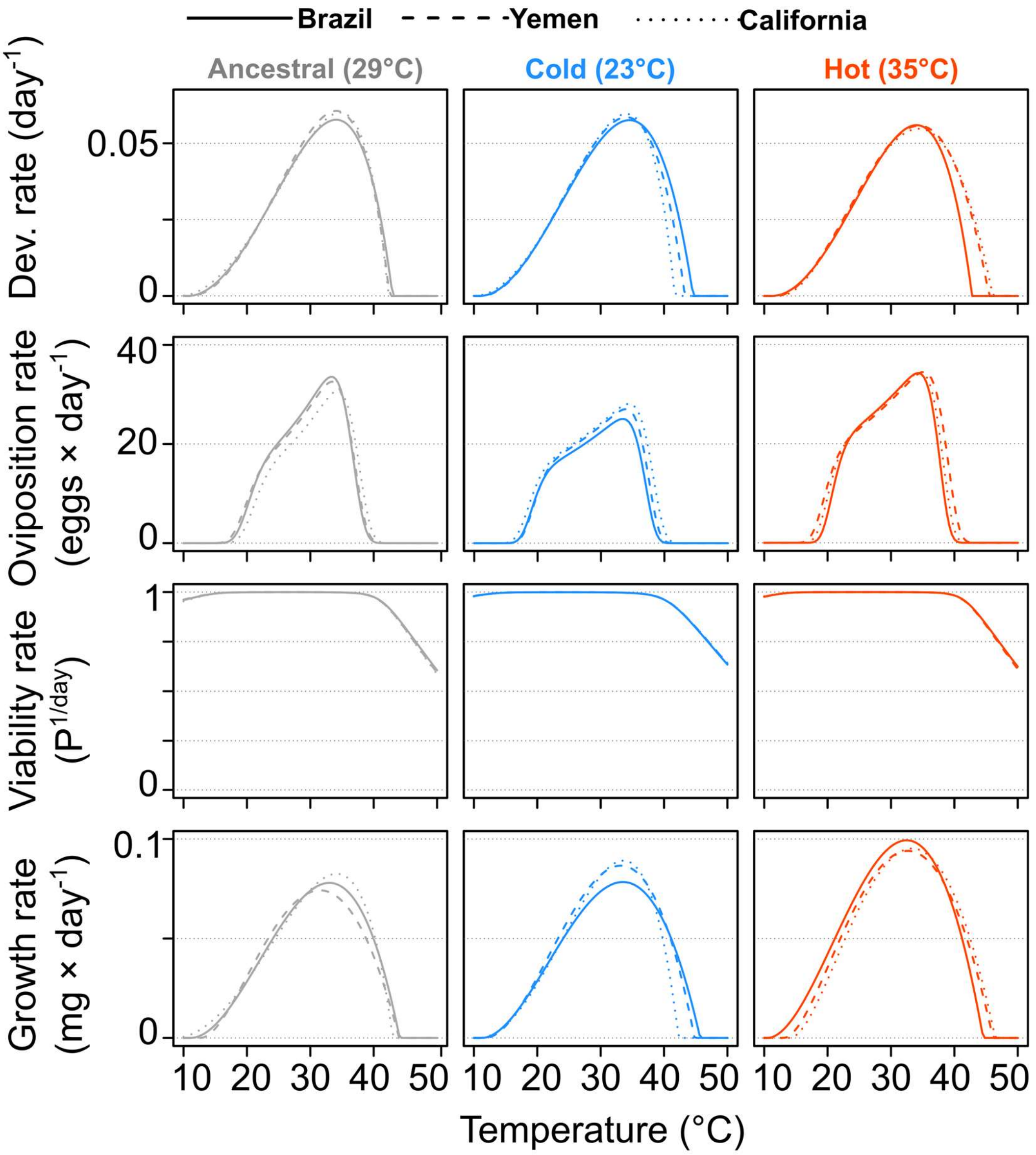

**Figure S5.** Estimated thermal performance curves for origin-specific traits expressed as daily rates.

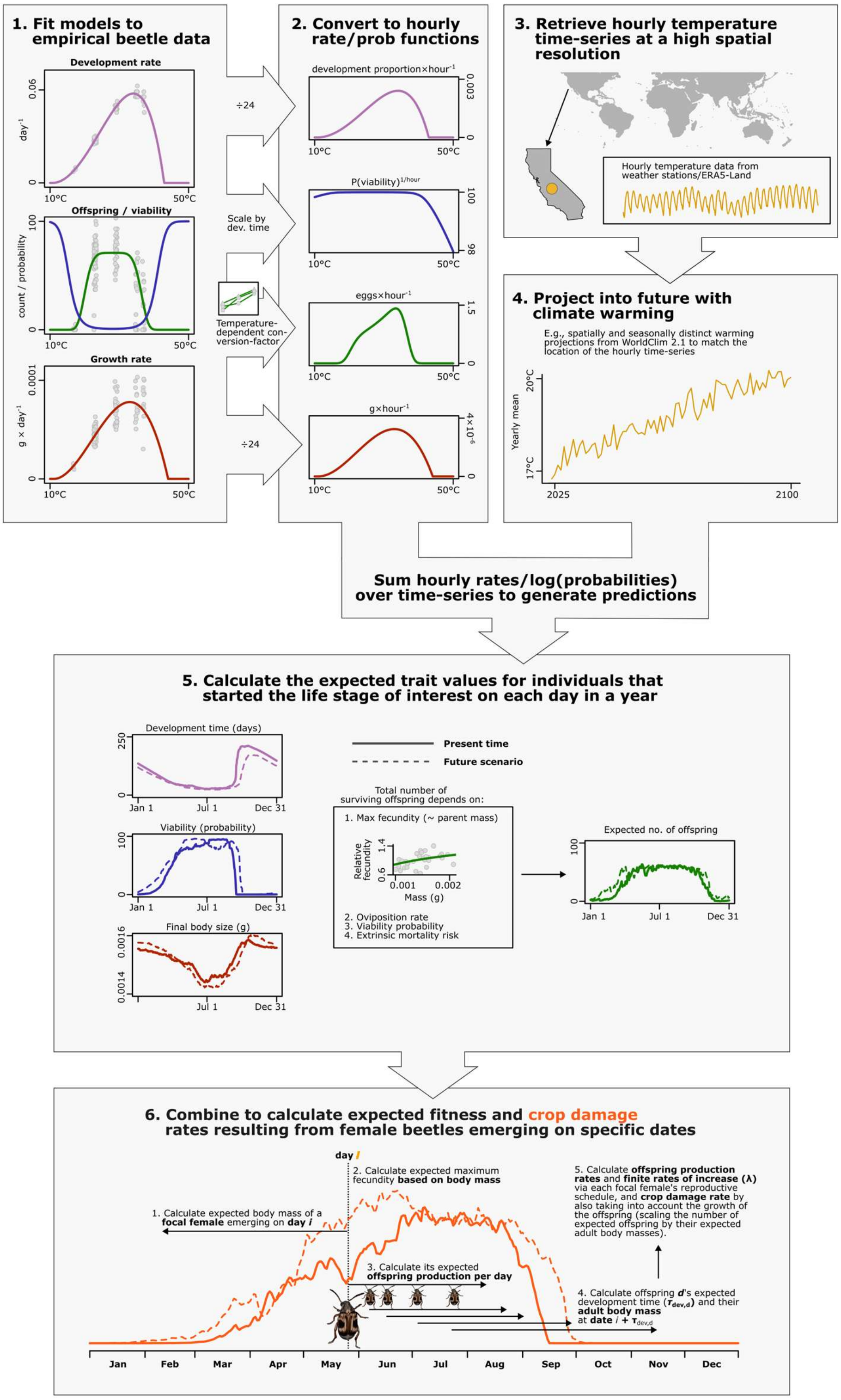

**Figure S6.** Overview of the process-based framework for projecting pest fitness and crop damage from empirically derived thermal reaction norms. (1) Bayesian thermal performance curves are fitted to empirical data for development rate, growth rate, and lifetime reproductive success (decomposed into viability and fecundity). (2) These are converted to instantaneous hourly functions, with viability expressed as an hourly probability and the other traits expressed as additive rates. (3) Hourly temperature time-series are retrieved at high spatial resolution and (4) projected into the different future warming scenarios. (5) Integrating the rate functions over these series gives the expected trait values of individuals initiating a given life stage on each day of the year. (6) Combining these trait values for focal females and their offspring yields a linear offspring production rate (short-term fitness proxy with interpretable units), the geometric finite rate of increase (λ; long-term niche suitability), and the crop damage rate, the last weighting offspring by their expected body mass (which scales with host consumption; Fig S7).

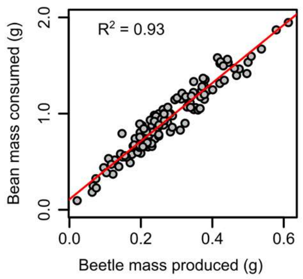

**Figure S7.** The relationship between beetle mass production and bean mass consumption, based on data from Burc et al. (2025).

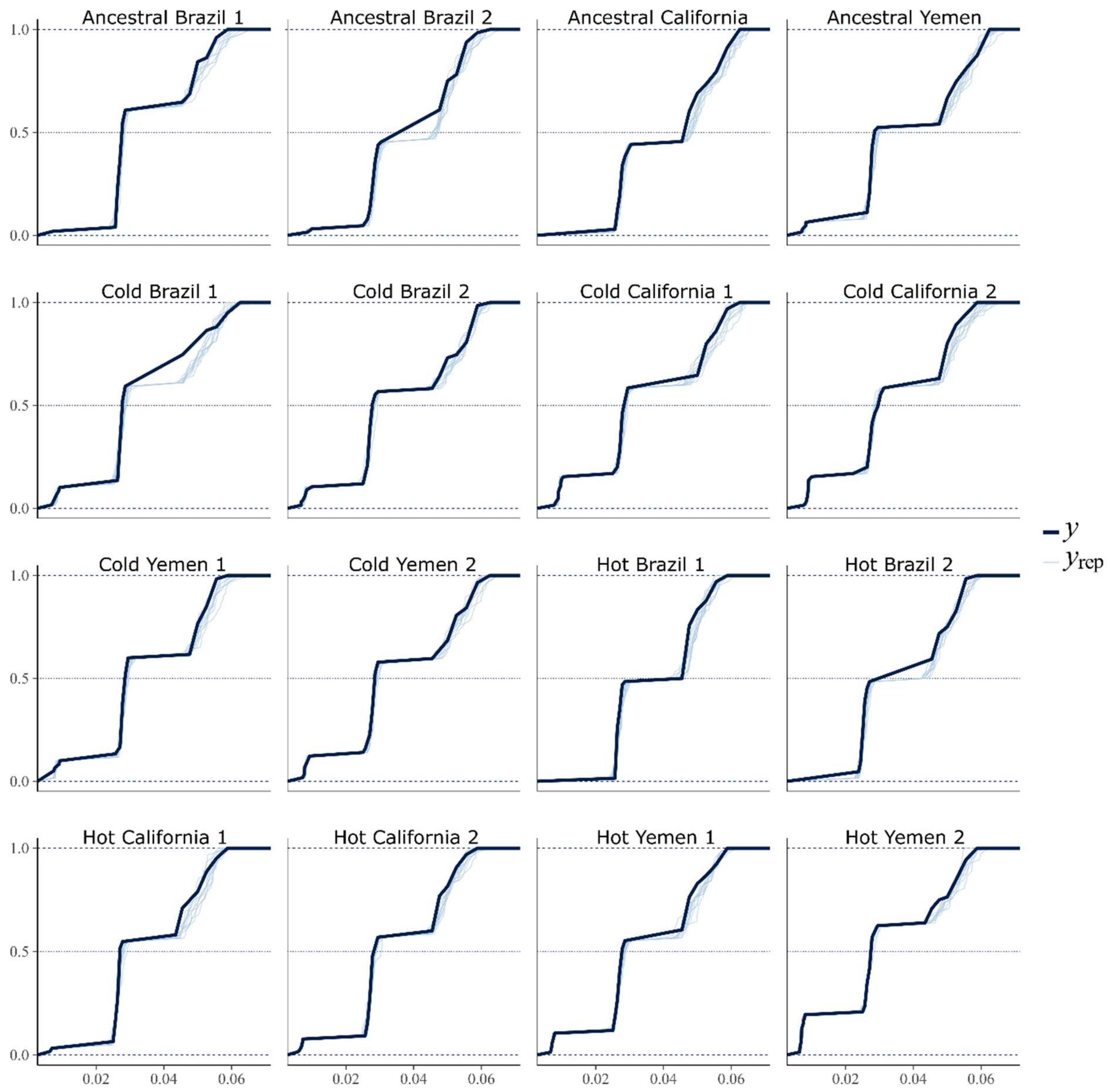

**Figure S8**. Posterior predictive checks for the origin-specific model of development rate thermal reaction norms. Panels show the empirical cumulative distributions for each replicate line. Thick black lines represent observed data; thin blue lines represent data simulated from the model (n = 10).

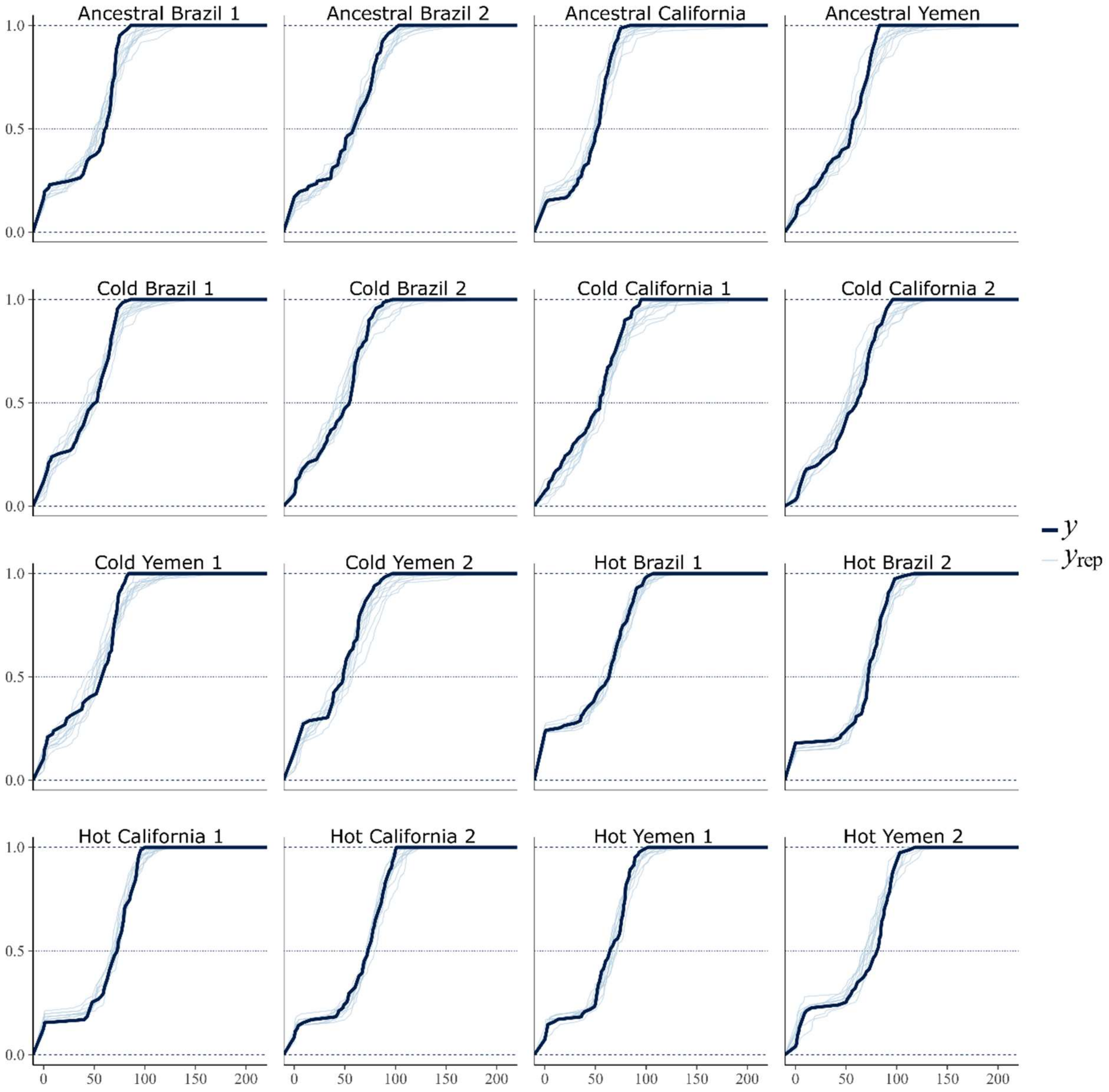

**Figure S9.** Posterior predictive checks for the origin-specific model of lifetime reproductive success thermal reaction norms. Panels show the empirical cumulative distributions for each replicate line. Thick black lines represent observed data; thin blue lines represent data simulated from the model (n = 10).

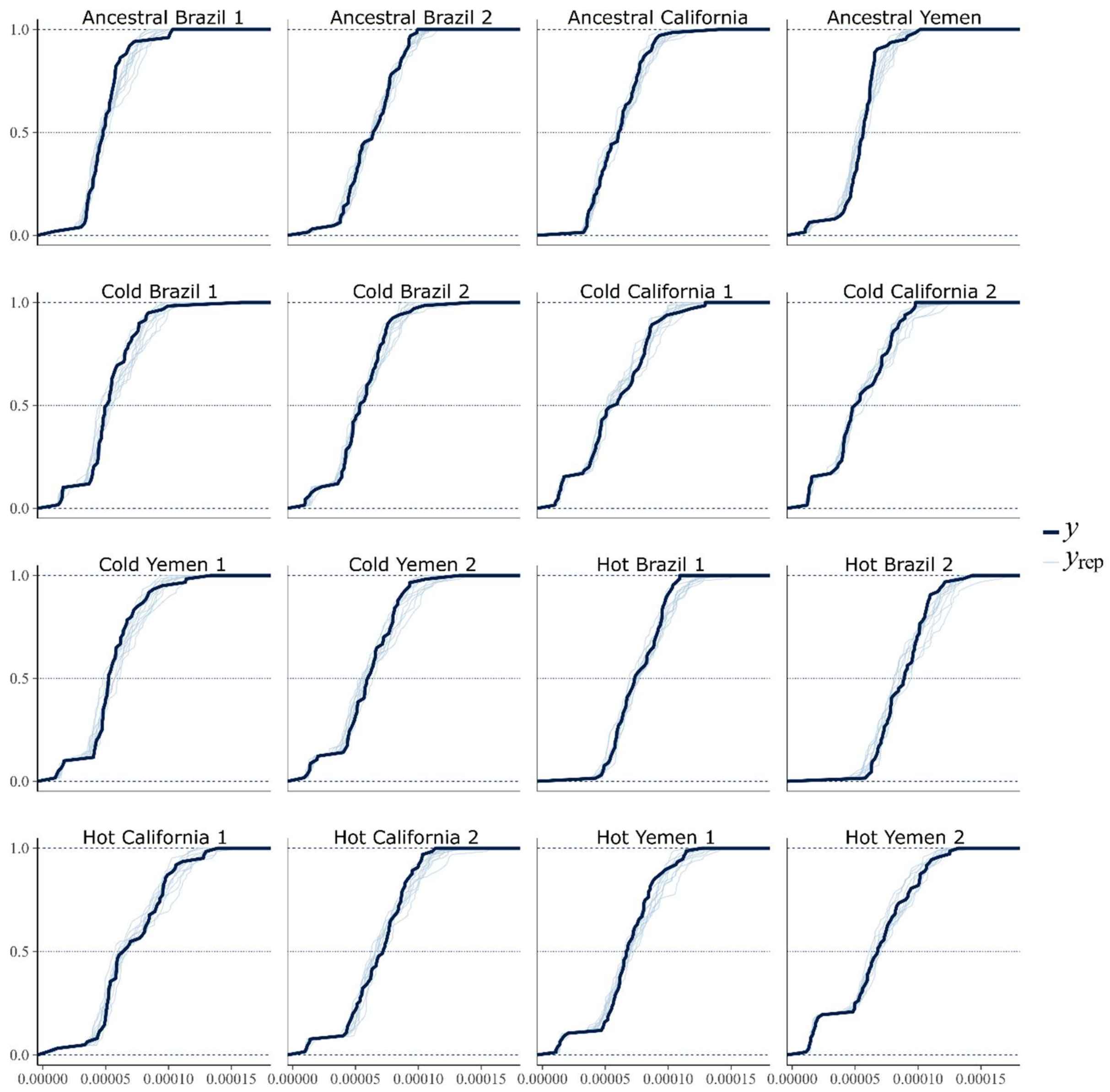

**Figure S10.** Posterior predictive checks for the origin-specific model of growth rate thermal reaction norms. Panels show empirical cumulative distributions for each replicate line. Thick black lines represent observed data; thin blue lines represent data simulated from the model (n = 10).

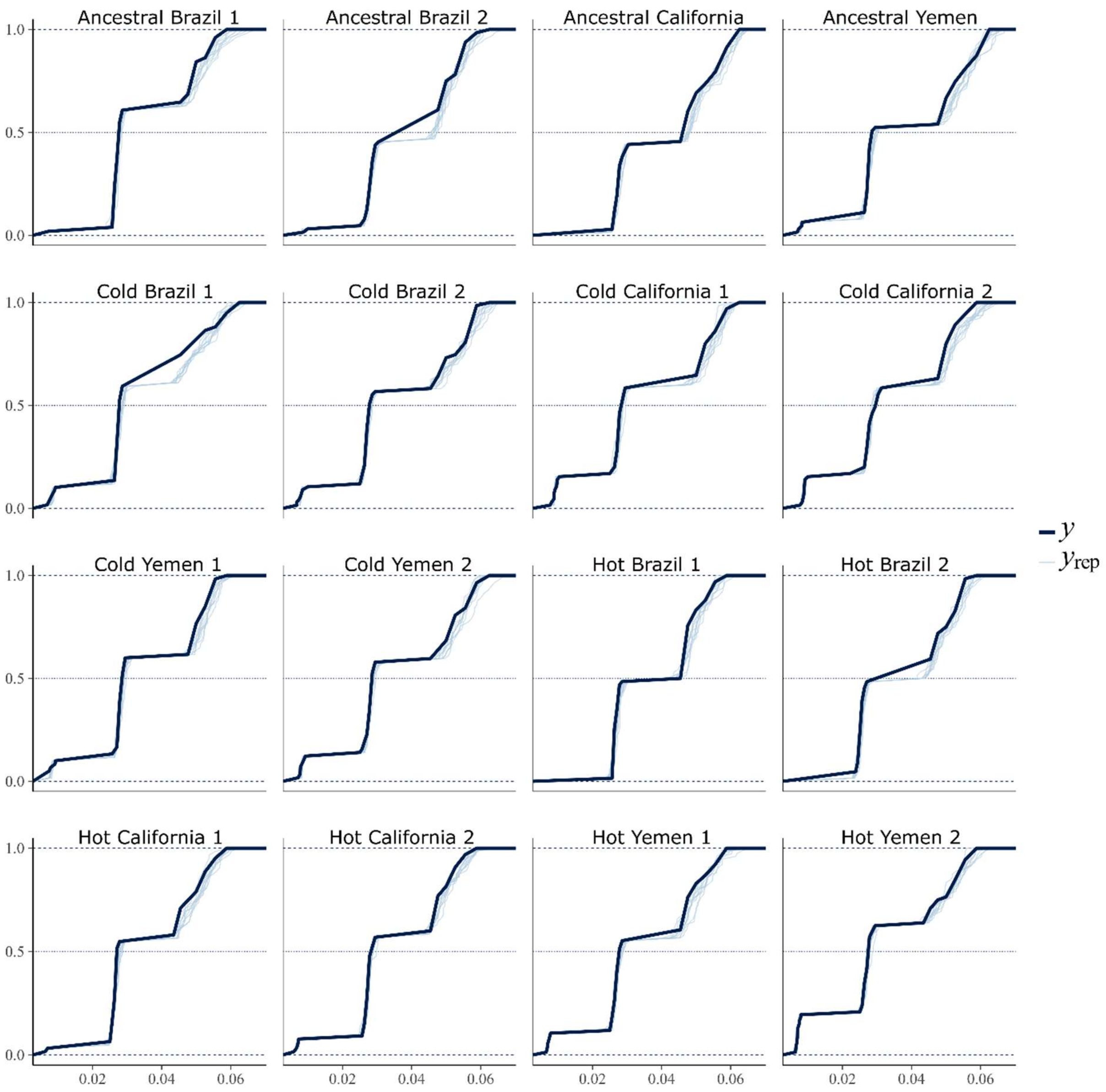

**Figure S11**. Posterior predictive checks for the “global” model of development rate thermal reaction norms. Panels show the empirical cumulative distributions for each replicate line. Thick black lines represent observed data; thin blue lines represent data simulated from the model (n = 10).

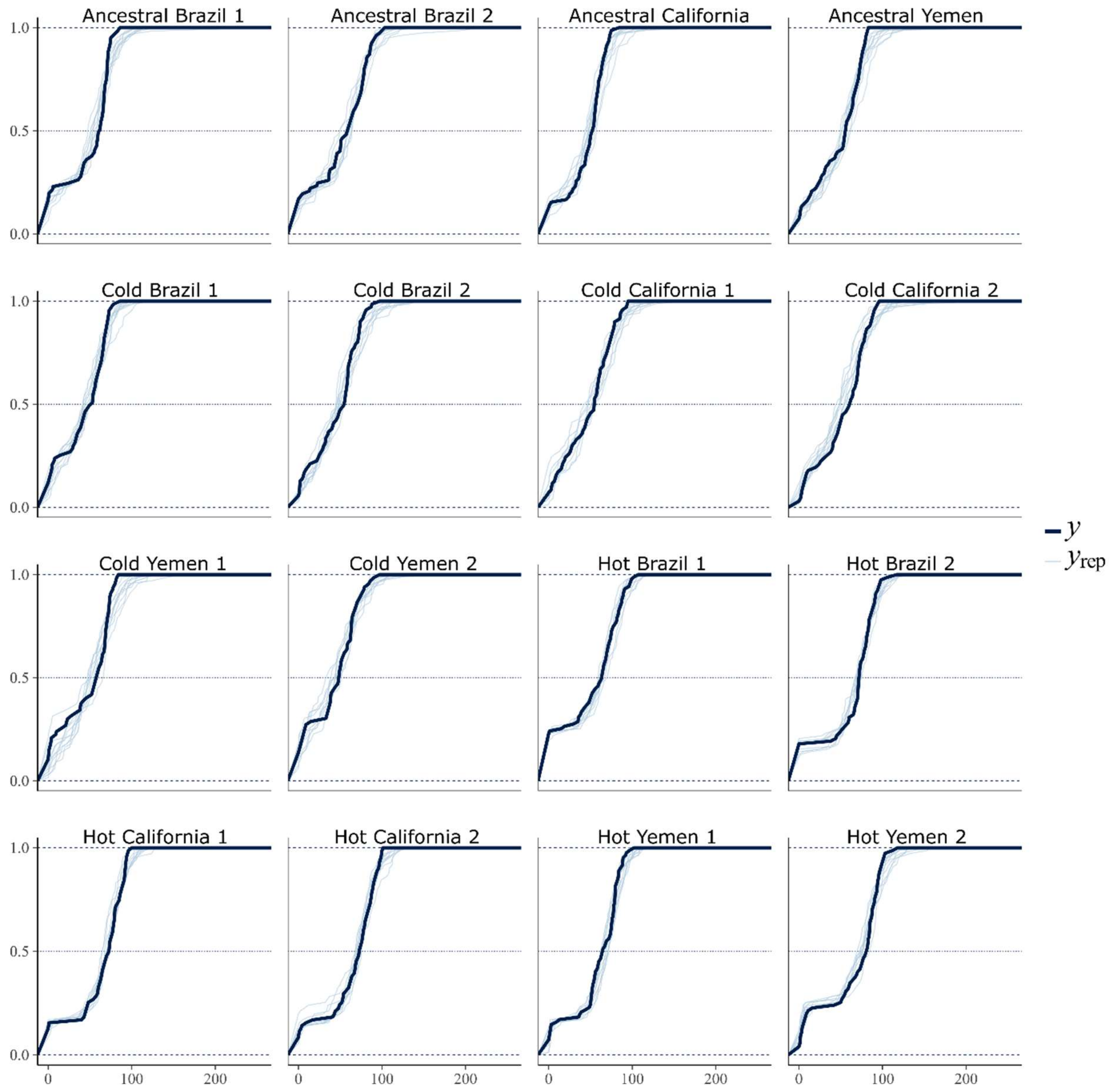

**Figure S12.** Posterior predictive checks for the "global" model of lifetime reproductive success thermal reaction norms. Panels show empirical cumulative distributions for each replicate line. Thick black lines represent observed data; thin blue lines represent data simulated from the model (n = 10).

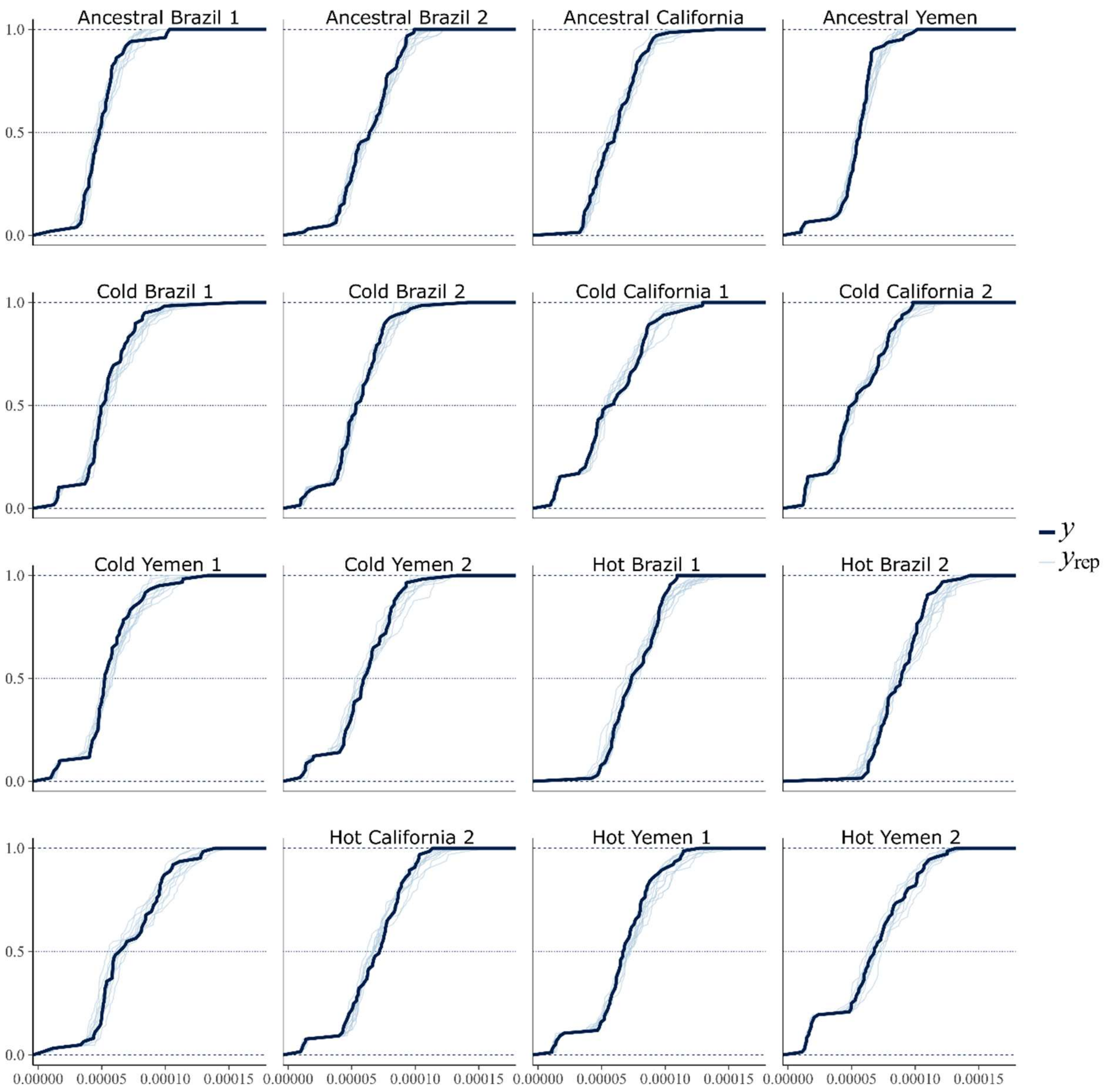

**Figure S13.** Posterior predictive checks for the “global” model of growth rate thermal reaction norms. Panels show empirical cumulative distributions for each replicate line. Thick black lines represent observed data; thin blue lines represent data simulated from the model (n = 10).

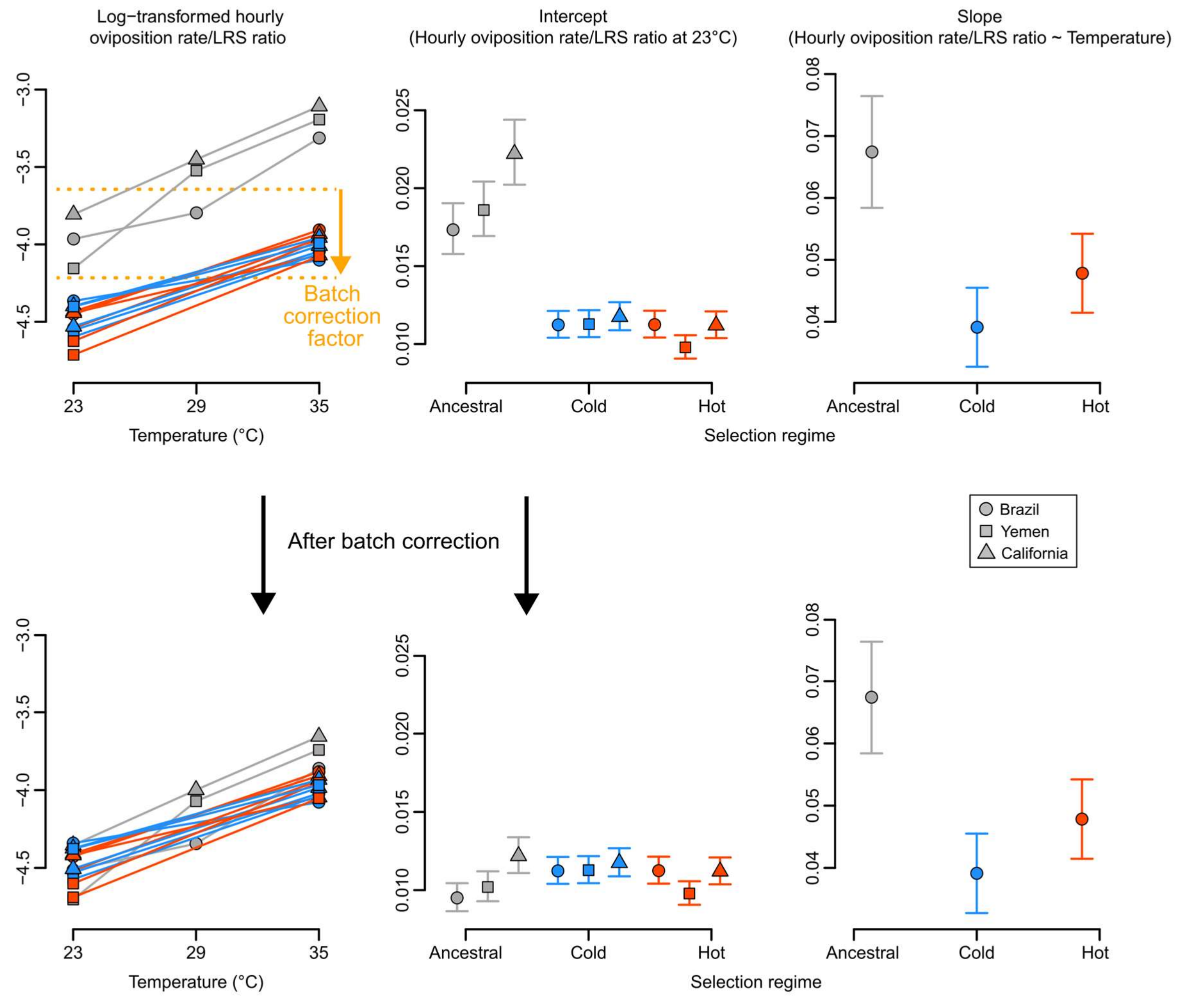

**Figure S14.** Visual demonstration of the conversion from lifetime reproductive success (LRS) to oviposition rate across temperatures. In the left panels, the relationship between oviposition rate and LRS are shown as ratios (y-axis) as predicted by temperature (x-axis). On the log-scale, the relationship between these ratios and temperature was approximately linear, allowing for a linear conversion of LRS reaction norms to oviposition rate reaction norms. Note that this relationship was scored in different experiments for the ancestral lines and the cold/hot adapted lines. However, another line was scored in both experimental batches, allowing for estimating a batch effect, and correcting the zero-centered intercept in the ancestral line to match the experimental batch of the cold/hot adapted lines (orange lines in the top left panel).

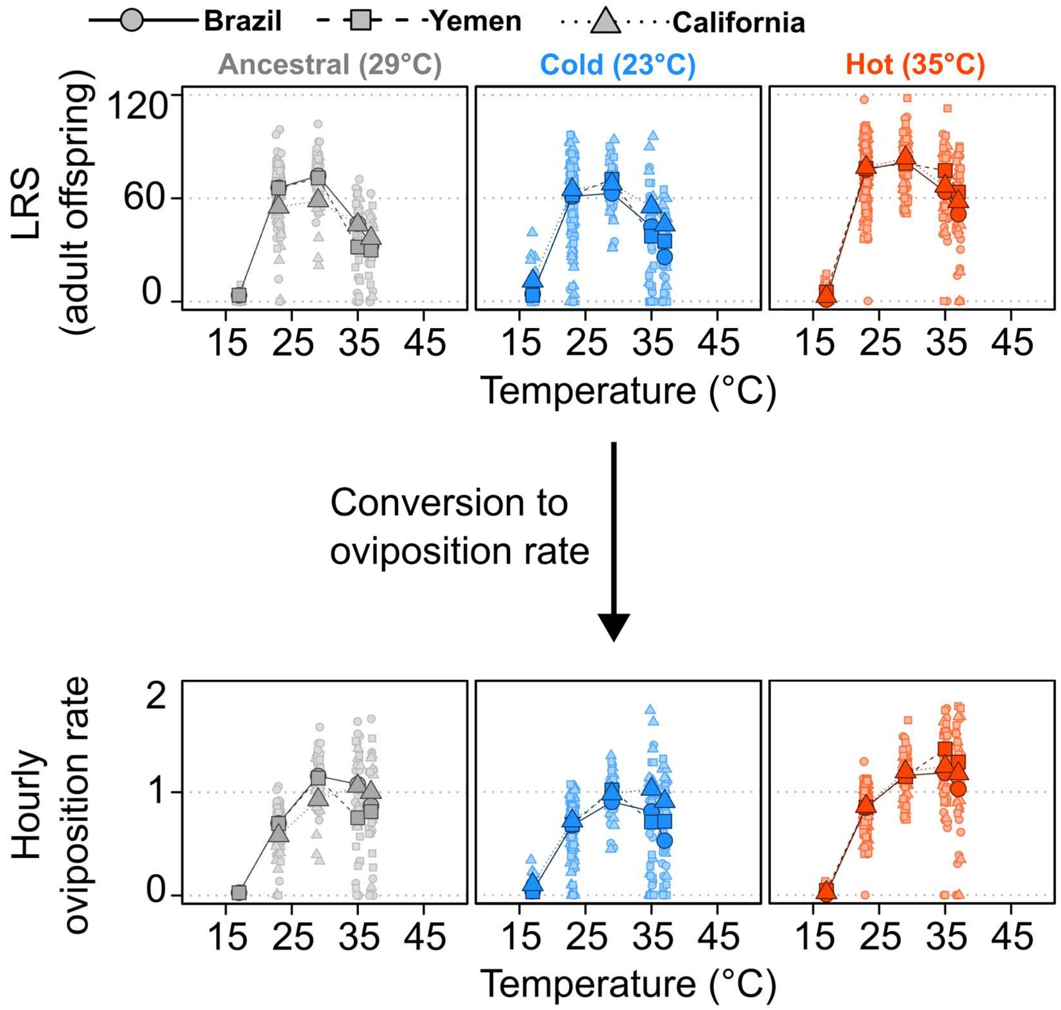

**Figure S15.** Data on lifetime reproductive success before and after conversion to oviposition rates.

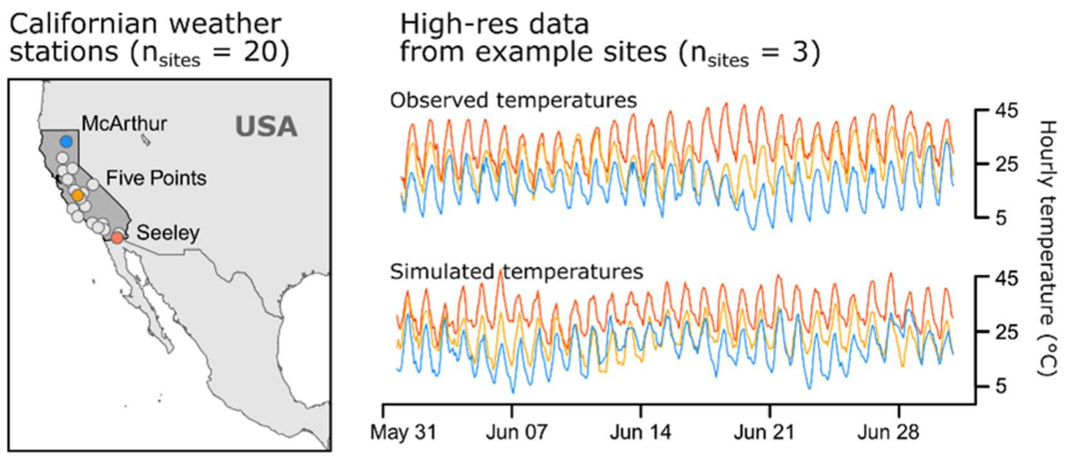

**Figure S16** The locations of the 20 Californian (USA) weather stations used to simulate temperature data, and a visual representation of both real and simulated hourly temperatures during summer for three example sites.

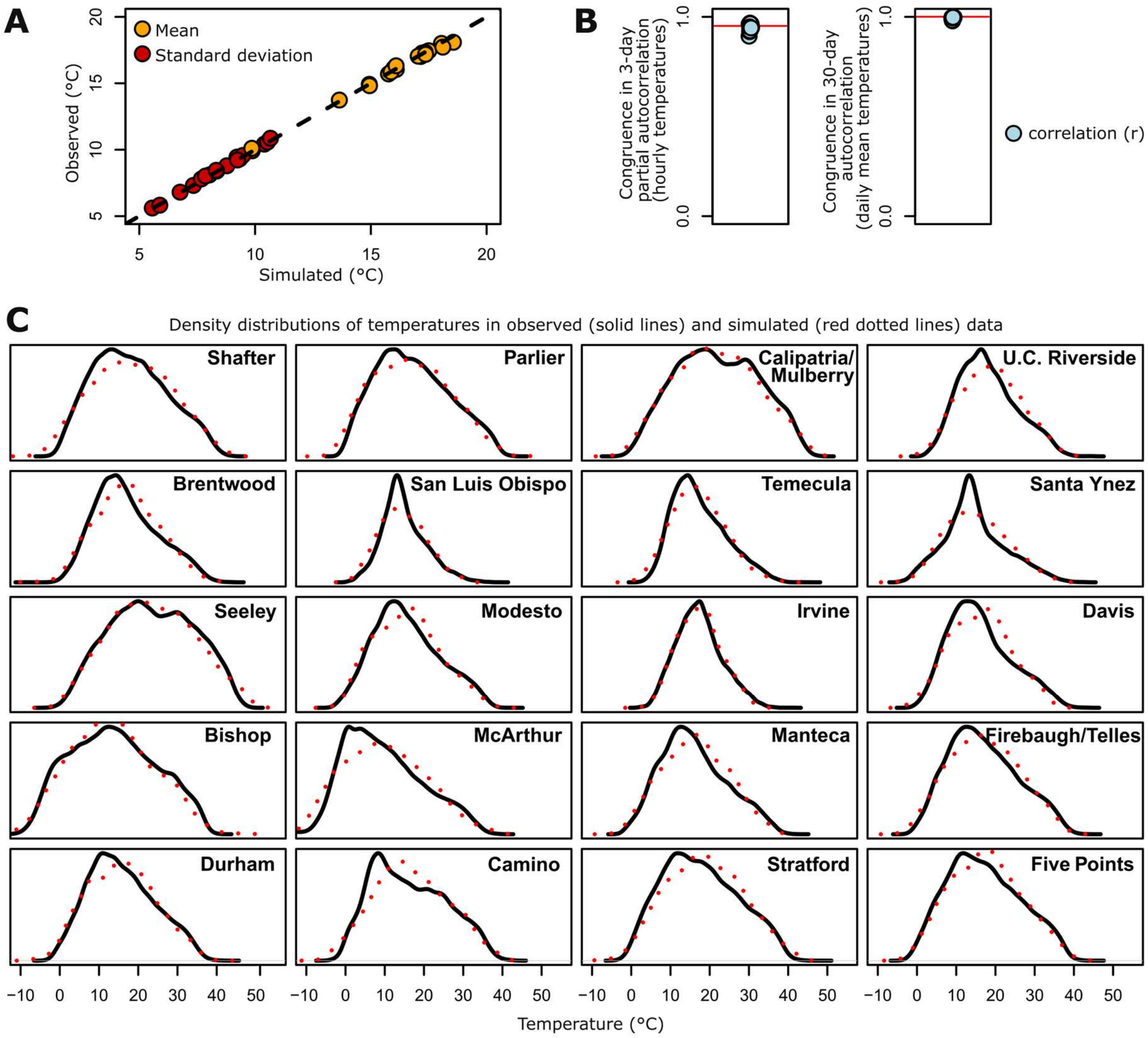

**Figure S17.** Overview of the similarities between observed and simulated temperatures for the different Californian (USA) weather station sites. (A) Correlations between mean temperatures (orange dots) and thermal variability (standard deviations, red dots) around those means. Each site is represented by one dot of each color and the dashed line represents a 1:1 relationship. (B) Summary of the correlation between observed and simulated data in partial autocorrelations in hourly temperatures over 3-days (see Fig. S20) and autocorrelations in daily average temperatures (see Fig. S21). Each site is one dot. (C) Density distributions of temperatures for the different sites. Black solid lines reflect real data and red dotted lines simulated data.

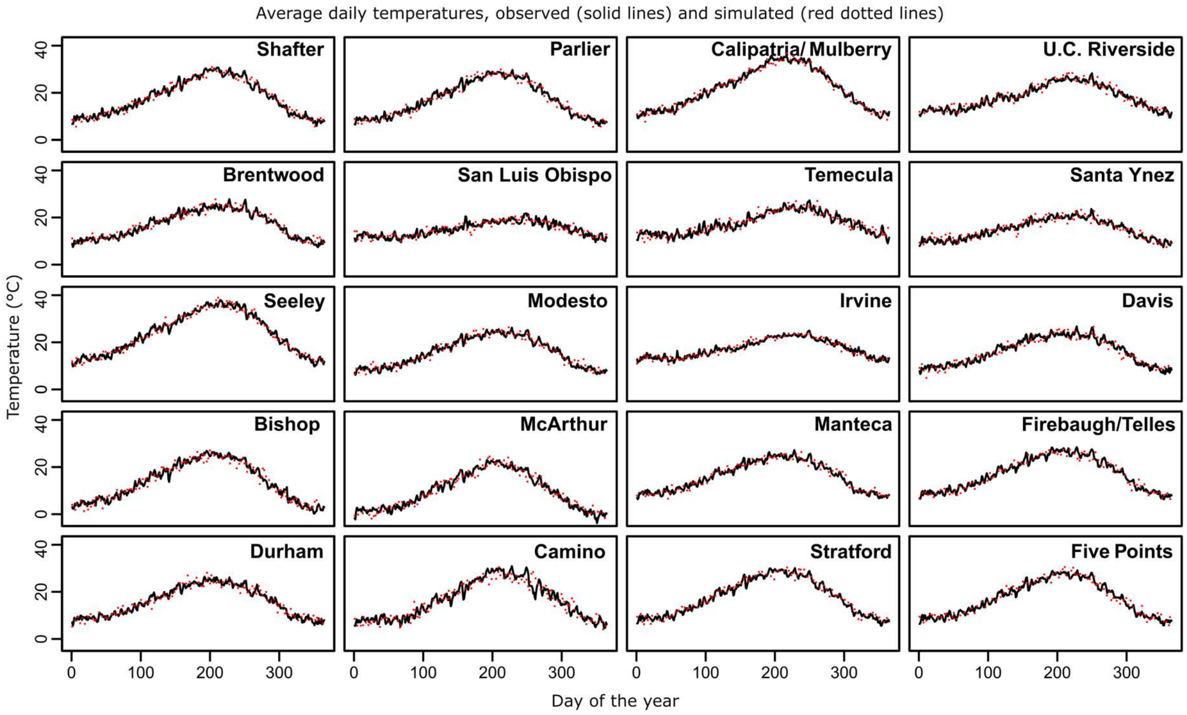

**Figure S18.** Average daily temperatures throughout the year for the different Californian (USA) weather station sites. Black solid lines reflect real data and red dotted lines simulated data. Their similarity shows the congruence in seasonality between observations and simulations.

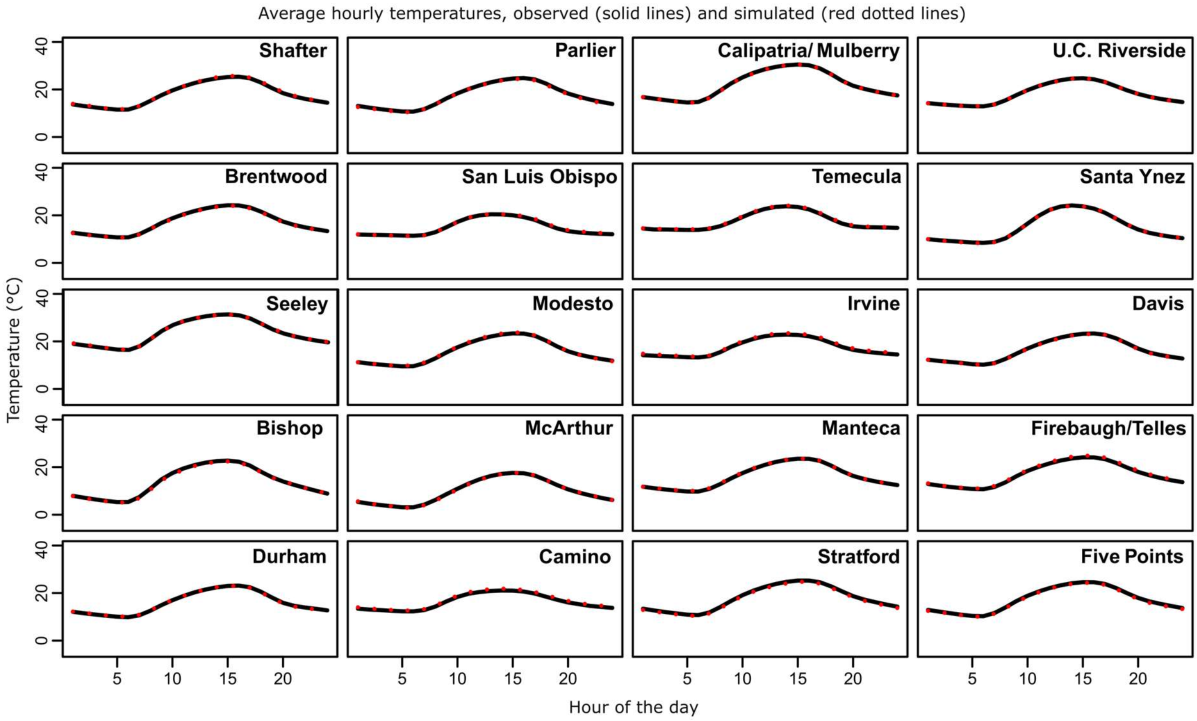

**Figure S19**. Average hourly temperatures throughout the day for the different Californian (USA) weather station sites. Black solid lines reflect real data and red dotted lines simulated data. Their similarity shows the congruence in average diurnal variability between observations and simulations.

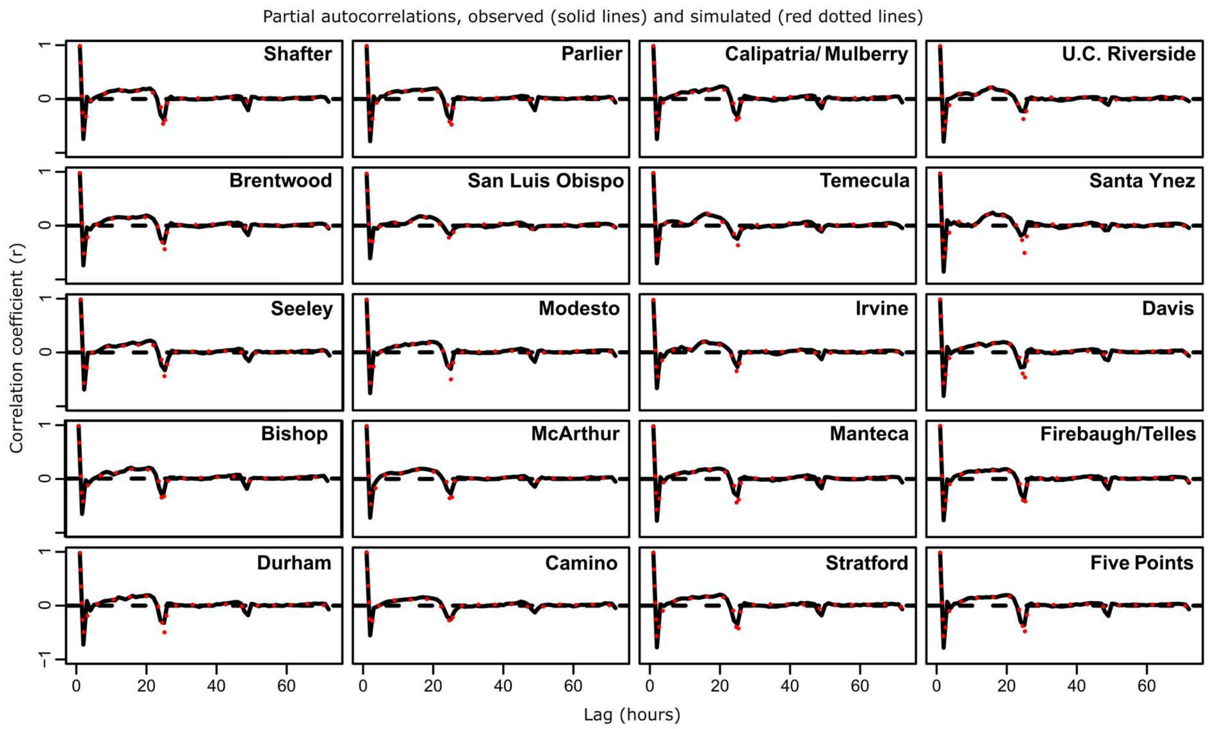

**Figure S20.** Hourly partial autocorrelations (i.e., discounting indirect correlations via consecutive autocorrelated time steps) within a lag of 3 days for each Californian (USA) weather station site. Black solid lines reflect real data and red dotted lines simulated data. Their similarity highlights a congruence in temporal structure between observations and simulations.

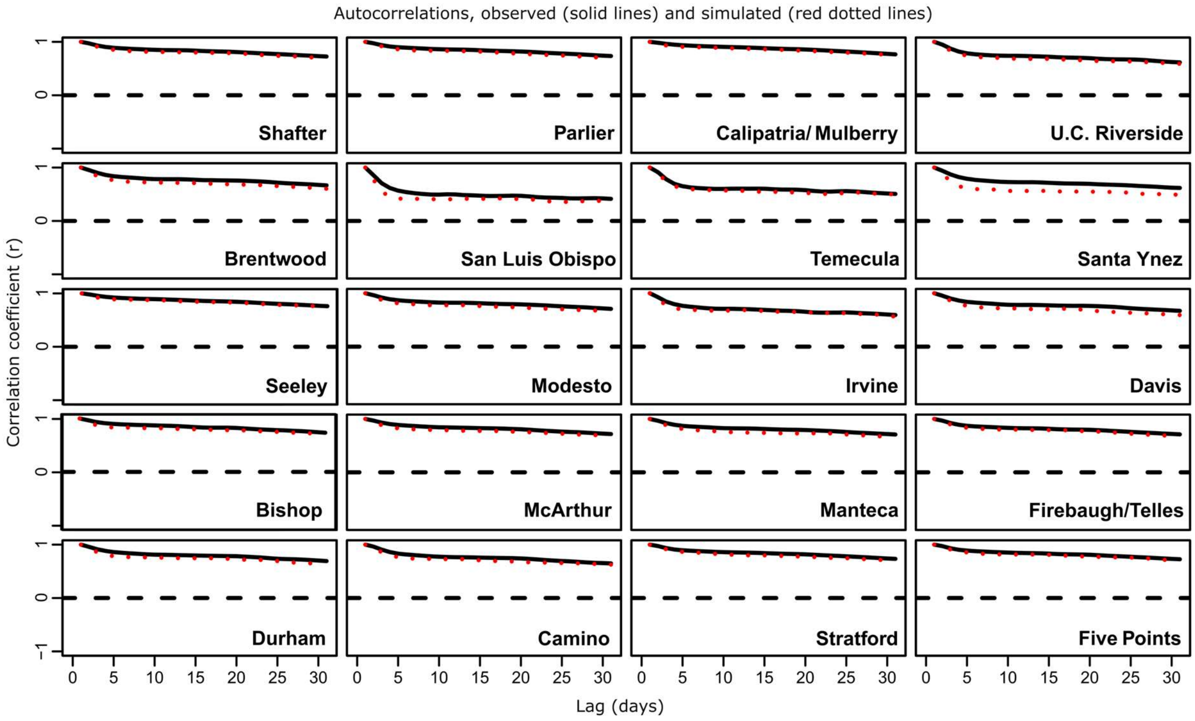

**Figure S21.** Daily autocorrelations within a lag phase of 30 days for each Californian (USA) weather station site. Black solid lines reflect real data and red dotted lines simulated data. Their similarity highlights a congruence in temporal structure between observations and simulations.

**Table S1.** Posterior parameter estimates (mode and 90% highest-density posterior interval) from the Bayesian thermal performance curve models for development rate, growth rate and offspring count, under origin-specific and global parameterizations. Origins (Brazil, USA, Yemen) and evolution regimes (ancestral, cold, hot) follow the experimental design. The priors used during fitting are listed in the final column. Rows with NA in the “Origin” column correspond to jointly estimated parameters.

| Model | Model type | Parameter | Origin | Evolution regime | Mode | 90% HDPI Lower | 90% HDPI Upper | Prior |
|---|---|---|---|---|---|---|---|---|
| Development rate | Origin-specific | Tmin | Brazil | Ancestral | 10.7 | 10.1 | 11.3 | normal(10, 4) |
| Development rate | Origin-specific | Tmin | USA | Ancestral | 9.31 | 7.54 | 11.8 | normal(10, 4) |
| Development rate | Origin-specific | Tmin | Yemen | Ancestral | 11.1 | 10.5 | 11.9 | normal(10, 4) |
| Development rate | Origin-specific | Tmin | Brazil | Cold | 11.0 | 10.4 | 11.4 | normal(10, 4) |
| Development rate | Origin-specific | Tmin | USA | Cold | 10.1 | 9.58 | 10.6 | normal(10, 4) |
| Development rate | Origin-specific | Tmin | Yemen | Cold | 11.2 | 10.7 | 11.6 | normal(10, 4) |
| Development rate | Origin-specific | Tmin | Brazil | Hot | 11.3 | 9.63 | 13.0 | normal(10, 4) |
| Development rate | Origin-specific | Tmin | USA | Hot | 12.4 | 11.8 | 12.9 | normal(10, 4) |
| Development rate | Origin-specific | Tmin | Yemen | Hot | 11.9 | 11.4 | 12.3 | normal(10, 4) |
| Development rate | Origin-specific | Tmax | Brazil | Ancestral | 43.0 | 41.5 | 44.6 | normal(45, 4) |
| Development rate | Origin-specific | Tmax | USA | Ancestral | 42.3 | 40.0 | 46.2 | normal(45, 4) |
| Development rate | Origin-specific | Tmax | Yemen | Ancestral | 42.5 | 40.7 | 44.4 | normal(45, 4) |
| Development rate | Origin-specific | Tmax | Brazil | Cold | 44.6 | 42.8 | 46.5 | normal(45, 4) |
| Development rate | Origin-specific | Tmax | USA | Cold | 41.7 | 40.6 | 43.2 | normal(45, 4) |
| Development rate | Origin-specific | Tmax | Yemen | Cold | 43.3 | 41.9 | 44.8 | normal(45, 4) |
| Development rate | Origin-specific | Tmax | Brazil | Hot | 42.8 | 41.1 | 45.2 | normal(45, 4) |
| Development rate | Origin-specific | Tmax | USA | Hot | 46.1 | 44.6 | 48.2 | normal(45, 4) |
| Development rate | Origin-specific | Tmax | Yemen | Hot | 45.5 | 43.8 | 47.2 | normal(45, 4) |
| Development rate | Origin-specific | Topt | Brazil | Ancestral | 34.1 | 33.5 | 34.6 | normal(35, 4) |
| Development rate | Origin-specific | Topt | USA | Ancestral | 34.7 | 34.1 | 35.6 | normal(35, 4) |
| Development rate | Origin-specific | Topt | Yemen | Ancestral | 34.1 | 33.4 | 34.9 | normal(35, 4) |
| Development rate | Origin-specific | Topt | Brazil | Cold | 34.5 | 34.0 | 35.2 | normal(35, 4) |
| Development rate | Origin-specific | Topt | USA | Cold | 34.1 | 33.6 | 34.5 | normal(35, 4) |
| Development rate | Origin-specific | Topt | Yemen | Cold | 34.1 | 33.5 | 34.5 | normal(35, 4) |
| Development rate | Origin-specific | Topt | Brazil | Hot | 34.0 | 33.5 | 34.6 | normal(35, 4) |
| Development rate | Origin-specific | Topt | USA | Hot | 34.4 | 33.9 | 35.1 | normal(35, 4) |
| Development rate | Origin-specific | Topt | Yemen | Hot | 34.4 | 33.9 | 35.1 | normal(35, 4) |
| Development rate | Origin-specific | rmax | Brazil | Ancestral | 0.0577 | 0.0561 | 0.0590 | normal(0.06, 0.06) |
| Development rate | Origin-specific | rmax | USA | Ancestral | 0.0601 | 0.0582 | 0.0628 | normal(0.06, 0.06) |
| Development rate | Origin-specific | rmax | Yemen | Ancestral | 0.0606 | 0.0585 | 0.0627 | normal(0.06, 0.06) |
| Development rate | Origin-specific | rmax | Brazil | Cold | 0.0576 | 0.0562 | 0.0589 | normal(0.06, 0.06) |
| Development rate | Origin-specific | rmax | USA | Cold | 0.0597 | 0.0582 | 0.0613 | normal(0.06, 0.06) |
| Development rate | Origin-specific | rmax | Yemen | Cold | 0.0586 | 0.0571 | 0.0602 | normal(0.06, 0.06) |
| Development rate | Origin-specific | rmax | Brazil | Hot | 0.0560 | 0.0543 | 0.0573 | normal(0.06, 0.06) |
| Development rate | Origin-specific | rmax | USA | Hot | 0.0548 | 0.0535 | 0.0562 | normal(0.06, 0.06) |
| Development rate | Origin-specific | rmax | Yemen | Hot | 0.0558 | 0.0546 | 0.0574 | normal(0.06, 0.06) |
| Development rate | Origin-specific | SD (Tmin) | NA | NA | 0.239 | 0.000312 | 0.577 | student_t(3, 0, 2.5) |
| Development rate | Origin-specific | SD (Tmax) | NA | NA | 0.728 | 0.00553 | 1.49 | student_t(3, 0, 2.5) |
| Development rate | Origin-specific | SD (Topt) | NA | NA | 0.312 | 0.150 | 0.634 | student_t(3, 0, 2.5) |
| Development rate | Origin-specific | SD (rmax) | NA | NA | 0.0134 | 0.00222 | 0.0283 | student_t(3, 0, 2.5) |
| Development rate | Origin-specific | Residual SD | NA | NA | 0.0462 | 0.0446 | 0.0481 | student_t(3, 0, 2.5) |
| Development rate | Global | Tmin | NA | Ancestral | 10.9 | 10.4 | 11.4 | normal(10, 4) |
| Development rate | Global | Tmin | NA | Cold | 10.8 | 10.4 | 11.2 | normal(10, 4) |
| Development rate | Global | Tmin | NA | Hot | 11.8 | 11.4 | 12.2 | normal(10, 4) |
| Development rate | Global | Tmax | NA | Ancestral | 43.1 | 41.6 | 45.4 | normal(45, 4) |
| Development rate | Global | Tmax | NA | Cold | 43.3 | 41.8 | 44.9 | normal(45, 4) |
| Development rate | Global | Tmax | NA | Hot | 44.3 | 42.9 | 46.6 | normal(45, 4) |
| Development rate | Global | Topt | NA | Ancestral | 34.2 | 33.9 | 34.6 | normal(35, 4) |
| Development rate | Global | Topt | NA | Cold | 34.2 | 33.9 | 34.5 | normal(35, 4) |
| Development rate | Global | Topt | NA | Hot | 34.2 | 33.9 | 34.7 | normal(35, 4) |
| Development rate | Global | rmax | NA | Ancestral | 0.0588 | 0.0571 | 0.0604 | normal(0.06, 0.06) |
| Development rate | Global | rmax | NA | Cold | 0.0580 | 0.0569 | 0.0596 | normal(0.06, 0.06) |
| Development rate | Global | rmax | NA | Hot | 0.0556 | 0.0545 | 0.0569 | normal(0.06, 0.06) |
| Development rate | Global | SD (replicate) | NA | NA | 0.0125 | 0.00101 | 0.0236 | student_t(3, 0, 2.5) |
| Development rate | Global | SD (replicate:temperature) | NA | NA | 0.0301 | 0.0244 | 0.0372 | student_t(3, 0, 2.5) |
| Development rate | Global | Residual SD | NA | NA | 0.0449 | 0.0434 | 0.0468 | student_t(3, 0, 2.5) |

**Table S1. Cont.**

| Model | Model type | Parameter | Origin | Evolution regime | Mode | 90% HDPI Lower | 90% HDPI Upper | Prior |
|---|---|---|---|---|---|---|---|---|
| Growth rate | Origin-specific | Tmin | Brazil | Ancestral | 11.1 | 9.96 | 12.3 | Flat |
| Growth rate | Origin-specific | Tmin | USA | Ancestral | 8.51 | 6.78 | 9.74 | Flat |
| Growth rate | Origin-specific | Tmin | Yemen | Ancestral | 12.6 | 11.3 | 13.8 | Flat |
| Growth rate | Origin-specific | Tmin | Brazil | Cold | 11.2 | 10.3 | 12.1 | Flat |
| Growth rate | Origin-specific | Tmin | USA | Cold | 10.7 | 9.75 | 11.4 | Flat |
| Growth rate | Origin-specific | Tmin | Yemen | Cold | 11.7 | 10.7 | 12.5 | Flat |
| Growth rate | Origin-specific | Tmin | Brazil | Hot | 10.5 | 8.81 | 11.9 | Flat |
| Growth rate | Origin-specific | Tmin | USA | Hot | 14.0 | 13.0 | 14.6 | Flat |
| Growth rate | Origin-specific | Tmin | Yemen | Hot | 13.0 | 12.1 | 13.7 | Flat |
| Growth rate | Origin-specific | Topt | Brazil | Ancestral | 33.0 | 32.3 | 33.7 | normal(35, 3) |
| Growth rate | Origin-specific | Topt | USA | Ancestral | 34.2 | 33.3 | 35.2 | normal(35, 3) |
| Growth rate | Origin-specific | Topt | Yemen | Ancestral | 31.5 | 30.8 | 32.5 | normal(35, 3) |
| Growth rate | Origin-specific | Topt | Brazil | Cold | 33.5 | 32.7 | 34.2 | normal(35, 3) |
| Growth rate | Origin-specific | Topt | USA | Cold | 33.6 | 33.1 | 34.3 | normal(35, 3) |
| Growth rate | Origin-specific | Topt | Yemen | Cold | 33.2 | 32.6 | 33.9 | normal(35, 3) |
| Growth rate | Origin-specific | Topt | Brazil | Hot | 32.5 | 31.8 | 33.2 | normal(35, 3) |
| Growth rate | Origin-specific | Topt | USA | Hot | 33.3 | 32.6 | 33.9 | normal(35, 3) |
| Growth rate | Origin-specific | Topt | Yemen | Hot | 32.6 | 32.0 | 33.3 | normal(35, 3) |
| Growth rate | Origin-specific | Tmax | Brazil | Ancestral | 44.1 | 42.5 | 45.3 | Flat |
| Growth rate | Origin-specific | Tmax | USA | Ancestral | 43.0 | 41.8 | 44.7 | Flat |
| Growth rate | Origin-specific | Tmax | Yemen | Ancestral | 44.0 | 42.7 | 45.2 | Flat |
| Growth rate | Origin-specific | Tmax | Brazil | Cold | 45.7 | 44.3 | 47.1 | Flat |
| Growth rate | Origin-specific | Tmax | USA | Cold | 42.3 | 41.1 | 43.7 | Flat |
| Growth rate | Origin-specific | Tmax | Yemen | Cold | 44.8 | 43.4 | 46.2 | Flat |
| Growth rate | Origin-specific | Tmax | Brazil | Hot | 44.5 | 43.2 | 45.8 | Flat |
| Growth rate | Origin-specific | Tmax | USA | Hot | 46.5 | 45.3 | 48.0 | Flat |
| Growth rate | Origin-specific | Tmax | Yemen | Hot | 46.1 | 44.8 | 47.6 | Flat |
| Growth rate | Origin-specific | rmax | Brazil | Ancestral | 0.0000780 | 0.0000708 | 0.0000857 | normal(0.00008, 0.00001) |
| Growth rate | Origin-specific | rmax | USA | Ancestral | 0.0000825 | 0.0000718 | 0.0000907 | normal(0.00008, 0.00001) |
| Growth rate | Origin-specific | rmax | Yemen | Ancestral | 0.0000742 | 0.0000675 | 0.0000865 | normal(0.00008, 0.00001) |
| Growth rate | Origin-specific | rmax | Brazil | Cold | 0.0000784 | 0.0000713 | 0.0000865 | normal(0.00008, 0.00001) |
| Growth rate | Origin-specific | rmax | USA | Cold | 0.0000891 | 0.0000821 | 0.0000982 | normal(0.00008, 0.00001) |
| Growth rate | Origin-specific | rmax | Yemen | Cold | 0.0000866 | 0.0000771 | 0.0000933 | normal(0.00008, 0.00001) |
| Growth rate | Origin-specific | rmax | Brazil | Hot | 0.0000994 | 0.0000879 | 0.000107 | normal(0.00008, 0.00001) |
| Growth rate | Origin-specific | rmax | USA | Hot | 0.0000952 | 0.0000851 | 0.000104 | normal(0.00008, 0.00001) |
| Growth rate | Origin-specific | rmax | Yemen | Hot | 0.0000940 | 0.0000847 | 0.000103 | normal(0.00008, 0.00001) |
| Growth rate | Origin-specific | SD (Tmin) | NA | NA | 0.389 | 0.00553 | 0.996 | normal(0, 0.25) |
| Growth rate | Origin-specific | SD (Topt) | NA | NA | 0.257 | 0.00101 | 0.606 | normal(0, 0.25) |
| Growth rate | Origin-specific | SD (Tmax) | NA | NA | 0.0593 | 0.000183 | 0.484 | normal(0, 0.25) |
| Growth rate | Origin-specific | SD (rmax) | NA | NA | 0.0641 | 0.0416 | 0.127 | normal(0, 0.25) |
| Growth rate | Origin-specific | Residual SD | NA | NA | 0.177 | 0.170 | 0.184 | student_t(3, 0, 2.5) |
| Growth rate | Global | Tmin | NA | Ancestral | 11.5 | 10.1 | 12.3 | Flat |
| Growth rate | Global | Tmin | NA | Cold | 10.9 | 10.0 | 11.7 | Flat |
| Growth rate | Global | Tmin | NA | Hot | 13.2 | 12.5 | 13.8 | Flat |
| Growth rate | Global | Topt | NA | Ancestral | 33.1 | 32.4 | 33.9 | normal(35, 3) |
| Growth rate | Global | Topt | NA | Cold | 33.7 | 33.1 | 34.5 | normal(35, 3) |
| Growth rate | Global | Topt | NA | Hot | 32.8 | 32.2 | 33.3 | normal(35, 3) |
| Growth rate | Global | Tmax | NA | Ancestral | 44.0 | 42.5 | 45.4 | Flat |
| Growth rate | Global | Tmax | NA | Cold | 44.2 | 42.8 | 45.7 | Flat |
| Growth rate | Global | Tmax | NA | Hot | 44.9 | 43.6 | 46.5 | Flat |
| Growth rate | Global | rmax | NA | Ancestral | 0.0000777 | 0.0000726 | 0.0000830 | normal(0.00008, 0.00001) |
| Growth rate | Global | rmax | NA | Cold | 0.0000847 | 0.0000800 | 0.0000892 | normal(0.00008, 0.00001) |
| Growth rate | Global | rmax | NA | Hot | 0.000101 | 0.0000952 | 0.000106 | normal(0.00008, 0.00001) |
| Growth rate | Global | SD (replicate) | NA | NA | 0.0470 | 0.0103 | 0.0862 | student_t(3, 0, 2.5) |
| Growth rate | Global | SD (replicate:temperature) | NA | NA | 0.0938 | 0.0738 | 0.116 | student_t(3, 0, 2.5) |
| Growth rate | Global | Residual SD | NA | NA | 0.171 | 0.164 | 0.177 | student_t(3, 0, 2.5) |

**Table S1. Cont.**

| Model | Model type | Parameter | Origin | Evolution regime | Mode | 90% HDPI Lower | Upper | Prior |
|---|---|---|---|---|---|---|---|---|
| Offspring count | Origin-specific | κ ancestral | NA | NA | 4.34 | 3.16 | 6.22 | lognormal((2.5), (2.5)) |
| Offspring count | Origin-specific | κ evolved | NA | NA | 5.48 | 4.84 | 6.34 | lognormal((2.5), (2.5)) |
| Offspring count | Origin-specific | a | NA | Ancestral | 0.0294 | 0.0220 | 0.0401 | lognormal(log(0.03), log(1.2)) |
| Offspring count | Origin-specific | a | NA | Cold | 0.0226 | 0.0180 | 0.0282 | lognormal(log(0.03), log(1.2)) |
| Offspring count | Origin-specific | a | NA | Hot | 0.0288 | 0.0215 | 0.0371 | lognormal(log(0.03), log(1.2)) |
| Offspring count | Origin-specific | Topt viability | NA | Ancestral | 28.1 | 26.3 | 29.5 | normal(28, 2) |
| Offspring count | Origin-specific | Topt viability | NA | Cold | 26.8 | 25.7 | 27.6 | normal(28, 2) |
| Offspring count | Origin-specific | Topt viability | NA | Hot | 27.4 | 25.6 | 28.8 | normal(28, 2) |
| Offspring count | Origin-specific | k | NA | Ancestral | -4.65 | -5.58 | -3.99 | normal(-4.5, 2) |
| Offspring count | Origin-specific | k | NA | Cold | -3.92 | -4.46 | -3.44 | normal(-4.5, 2) |
| Offspring count | Origin-specific | k | NA | Hot | -5.48 | -6.68 | -4.79 | normal(-4.5, 2) |
| Offspring count | Origin-specific | Topt fecundity | Brazil | Ancestral | 28.1 | 27.3 | 28.9 | normal(29, 5) |
| Offspring count | Origin-specific | Topt fecundity | USA | Ancestral | 29.0 | 28.0 | 30.2 | normal(29, 5) |
| Offspring count | Origin-specific | Topt fecundity | Yemen | Ancestral | 28.0 | 26.6 | 29.0 | normal(29, 5) |
| Offspring count | Origin-specific | Topt fecundity | Brazil | Cold | 27.9 | 27.3 | 28.5 | normal(29, 5) |
| Offspring count | Origin-specific | Topt fecundity | USA | Cold | 28.2 | 27.5 | 28.9 | normal(29, 5) |
| Offspring count | Origin-specific | Topt fecundity | Yemen | Cold | 28.2 | 27.6 | 28.9 | normal(29, 5) |
| Offspring count | Origin-specific | Topt fecundity | Brazil | Hot | 29.0 | 28.5 | 29.7 | normal(29, 5) |
| Offspring count | Origin-specific | Topt fecundity | USA | Hot | 28.9 | 28.4 | 29.6 | normal(29, 5) |
| Offspring count | Origin-specific | Topt fecundity | Yemen | Hot | 29.0 | 28.3 | 29.7 | normal(29, 5) |
| Offspring count | Origin-specific | fecmax | Brazil | Ancestral | 70.3 | 65.1 | 77.3 | normal(70, 25) |
| Offspring count | Origin-specific | fecmax | USA | Ancestral | 60.4 | 51.8 | 68.0 | normal(70, 25) |
| Offspring count | Origin-specific | fecmax | Yemen | Ancestral | 68.0 | 60.4 | 77.6 | normal(70, 25) |
| Offspring count | Origin-specific | fecmax | Brazil | Cold | 62.2 | 56.5 | 67.9 | normal(70, 25) |
| Offspring count | Origin-specific | fecmax | USA | Cold | 67.3 | 61.2 | 72.8 | normal(70, 25) |
| Offspring count | Origin-specific | fecmax | Yemen | Cold | 65.8 | 60.1 | 72.2 | normal(70, 25) |
| Offspring count | Origin-specific | fecmax | Brazil | Hot | 81.8 | 75.1 | 86.8 | normal(70, 25) |
| Offspring count | Origin-specific | fecmax | USA | Hot | 81.3 | 75.9 | 86.9 | normal(70, 25) |
| Offspring count | Origin-specific | fecmax | Yemen | Hot | 79.8 | 74.4 | 85.5 | normal(70, 25) |
| Offspring count | Origin-specific | breadth | Brazil | Ancestral | 21.3 | 20.1 | 23.3 | lognormal(log(22), log(1.1)) |
| Offspring count | Origin-specific | breadth | USA | Ancestral | 21.5 | 19.7 | 24.9 | lognormal(log(22), log(1.1)) |
| Offspring count | Origin-specific | breadth | Yemen | Ancestral | 22.2 | 20.6 | 25.0 | lognormal(log(22), log(1.1)) |
| Offspring count | Origin-specific | breadth | Brazil | Cold | 22.0 | 21.2 | 22.9 | lognormal(log(22), log(1.1)) |
| Offspring count | Origin-specific | breadth | USA | Cold | 24.1 | 23.0 | 25.4 | lognormal(log(22), log(1.1)) |
| Offspring count | Origin-specific | breadth | Yemen | Cold | 22.3 | 21.5 | 23.3 | lognormal(log(22), log(1.1)) |
| Offspring count | Origin-specific | breadth | Brazil | Hot | 20.9 | 20.0 | 21.8 | lognormal(log(22), log(1.1)) |
| Offspring count | Origin-specific | breadth | USA | Hot | 22.3 | 21.5 | 23.3 | lognormal(log(22), log(1.1)) |
| Offspring count | Origin-specific | breadth | Yemen | Hot | 23.9 | 22.9 | 25.1 | lognormal(log(22), log(1.1)) |
| Offspring count | Origin-specific | shape (17°C) | NA | Ancestral | -1.88 | -3.18 | -0.139 | Flat |
| Offspring count | Origin-specific | shape (23°C) | NA | Ancestral | 2.82 | 2.54 | 3.16 | Flat |
| Offspring count | Origin-specific | shape (29°C) | NA | Ancestral | 3.34 | 2.93 | 3.86 | Flat |
| Offspring count | Origin-specific | shape (35°C) | NA | Ancestral | 1.05 | 0.568 | 1.50 | Flat |
| Offspring count | Origin-specific | shape (37°C) | NA | Ancestral | 1.21 | 0.648 | 1.81 | Flat |
| Offspring count | Origin-specific | shape (17°C) | NA | Cold | 0.693 | 0.170 | 1.25 | Flat |
| Offspring count | Origin-specific | shape (23°C) | NA | Cold | 2.31 | 2.07 | 2.51 | Flat |
| Offspring count | Origin-specific | shape (29°C) | NA | Cold | 3.59 | 3.09 | 4.18 | Flat |
| Offspring count | Origin-specific | shape (35°C) | NA | Cold | 1.14 | 0.772 | 1.52 | Flat |
| Offspring count | Origin-specific | shape (37°C) | NA | Cold | 1.50 | 1.06 | 1.87 | Flat |
| Offspring count | Origin-specific | shape (17°C) | NA | Hot | -0.126 | -0.769 | 0.629 | Flat |
| Offspring count | Origin-specific | shape (23°C) | NA | Hot | 3.54 | 3.28 | 3.77 | Flat |
| Offspring count | Origin-specific | shape (29°C) | NA | Hot | 4.08 | 3.71 | 4.67 | Flat |
| Offspring count | Origin-specific | shape (35°C) | NA | Hot | 2.95 | 2.50 | 3.41 | Flat |
| Offspring count | Origin-specific | shape (37°C) | NA | Hot | 2.75 | 2.29 | 3.20 | Flat |
| Offspring count | Origin-specific | SD (Topt) | NA | NA | 0.343 | 0.144 | 0.748 | student_t(3, 0, 2.5) |
| Offspring count | Origin-specific | SD (fecmax) | NA | NA | 3.43 | 1.84 | 6.17 | student_t(3, 0, 2.5) |
| Offspring count | Origin-specific | SD (breadth) | NA | NA | 0.384 | 0.00290 | 1.04 | student_t(3, 0, 2.5) |

### Table S1. Cont.

| Model | Model type | Parameter | Origin | Evolution regime | Mode | 90% HDPI Lower | 90% HDPI Upper | Prior |
|---|---|---|---|---|---|---|---|---|
| Offspring count | Global | Topt | NA | Ancestral | 28.4 | 27.9 | 28.8 | normal(29, 5) |
| Offspring count | Global | Topt | NA | Cold | 28.3 | 28.0 | 28.6 | normal(29, 5) |
| Offspring count | Global | Topt | NA | Hot | 28.8 | 28.3 | 29.2 | normal(29, 5) |
| Offspring count | Global | fecmax | NA | Ancestral | 67.3 | 61.9 | 73.7 | normal(70, 25) |
| Offspring count | Global | fecmax | NA | Cold | 67.0 | 62.2 | 72.3 | normal(70, 25) |
| Offspring count | Global | fecmax | NA | Hot | 79.4 | 74.8 | 85.4 | normal(70, 25) |
| Offspring count | Global | breadth | NA | Ancestral | 21.8 | 20.9 | 24.1 | lognormal(log(22), log(1.1)) |
| Offspring count | Global | breadth | NA | Cold | 24.0 | 22.9 | 25.1 | lognormal(log(22), log(1.1)) |
| Offspring count | Global | breadth | NA | Hot | 23.4 | 22.4 | 24.2 | lognormal(log(22), log(1.1)) |
| Offspring count | Global | a | NA | Ancestral | 0.0307 | 0.0226 | 0.0413 | lognormal(log(0.03), log(1.2)) |
| Offspring count | Global | a | NA | Cold | 0.0254 | 0.0200 | 0.0341 | lognormal(log(0.03), log(1.2)) |
| Offspring count | Global | a | NA | Hot | 0.0336 | 0.0254 | 0.0447 | lognormal(log(0.03), log(1.2)) |
| Offspring count | Global | Topt viability | NA | Ancestral | 28.7 | 26.4 | 30.7 | normal(28, 2) |
| Offspring count | Global | Topt viability | NA | Cold | 26.8 | 25.3 | 28.4 | normal(28, 2) |
| Offspring count | Global | Topt viability | NA | Hot | 29.0 | 27.8 | 30.5 | normal(28, 2) |
| Offspring count | Global | k | NA | Ancestral | -5.17 | -6.60 | -4.04 | normal(-4.5, 2) |
| Offspring count | Global | k | NA | Cold | -4.76 | -5.93 | -3.91 | normal(-4.5, 2) |
| Offspring count | Global | k | NA | Hot | -5.76 | -7.04 | -4.80 | normal(-4.5, 2) |
| Offspring count | Global | κ | NA | Ancestral | 3.78 | 2.58 | 5.68 | lognormal((2.5), (2.5)) |
| Offspring count | Global | κ | NA | Cold | 3.32 | 2.50 | 5.00 | lognormal((2.5), (2.5)) |
| Offspring count | Global | κ | NA | Hot | 4.60 | 3.65 | 7.43 | lognormal((2.5), (2.5)) |
| Offspring count | Global | shape (17°C) | NA | Ancestral | -1.82 | -3.14 | 0.347 | Flat |
| Offspring count | Global | shape (23°C) | NA | Ancestral | 2.85 | 2.52 | 3.13 | Flat |
| Offspring count | Global | shape (29°C) | NA | Ancestral | 3.46 | 2.96 | 3.94 | Flat |
| Offspring count | Global | shape (35°C) | NA | Ancestral | 1.03 | 0.648 | 1.56 | Flat |
| Offspring count | Global | shape (37°C) | NA | Ancestral | 1.02 | 0.467 | 1.52 | Flat |
| Offspring count | Global | shape (17°C) | NA | Cold | 0.0708 | -0.362 | 0.580 | Flat |
| Offspring count | Global | shape (23°C) | NA | Cold | 2.32 | 2.09 | 2.53 | Flat |
| Offspring count | Global | shape (29°C) | NA | Cold | 3.65 | 3.15 | 4.23 | Flat |
| Offspring count | Global | shape (35°C) | NA | Cold | 1.20 | 0.841 | 1.62 | Flat |
| Offspring count | Global | shape (37°C) | NA | Cold | 1.40 | 1.03 | 1.81 | Flat |
| Offspring count | Global | shape (17°C) | NA | Hot | -0.111 | -0.855 | 0.567 | Flat |
| Offspring count | Global | shape (23°C) | NA | Hot | 3.52 | 3.28 | 3.78 | Flat |
| Offspring count | Global | shape (29°C) | NA | Hot | 4.19 | 3.71 | 4.71 | Flat |
| Offspring count | Global | shape (35°C) | NA | Hot | 3.09 | 2.59 | 3.58 | Flat |
| Offspring count | Global | shape (37°C) | NA | Hot | 2.56 | 2.13 | 3.02 | Flat |
| Offspring count | Global | SD (replicate) viability | NA | NA | 0.381 | 0.000123 | 1.21 | student_t(3, 0, 2.5) |
| Offspring count | Global | SD (replicate:temperature) viability | NA | NA | 1.51 | 1.01 | 2.19 | student_t(3, 0, 2.5) |
| Offspring count | Global | SD (replicate) fecundity | NA | NA | 0.0679 | 0.0363 | 0.111 | student_t(3, 0, 2.5) |
| Offspring count | Global | SD (replicate:temperature) fecundity | NA | NA | 0.0531 | 0.00404 | 0.0872 | student_t(3, 0, 2.5) |

**Table S2.** Predicted changes in fitness (per-capita offspring production rate; offspring × day$^{-1}$) under different warming scenarios across 20 Californian (USA) sites by the year 2100. The top row shows changes via strictly plastic effects in the ancestral lines (grey), serving as a baseline of ~30% increase in median offspring production rates per °C of warming. The following rows represent additive deviations from this baseline, representing additional fitness changes introduced by evolution (hot = red, cold = blue). Responses have been calculated for the full phenotypes (i.e., by integrating all four evolved reaction norms; trait = all). To investigate trait-specific contributions, responses have also been calculated separately for each trait (i.e., all other reaction norms fixed to their ancestral states). Boxplots show medians (denoted in bold), interquartile ranges, and full ranges (denoted in brackets; arrows denote ranges exceeding the plots' x-limit) across the sites. Evolutionary deviations (colored) from the ancestral baseline (grey) for all warming rates are displayed in the right column.

| | Selection regime | Trait | Warming rate 0°C/y | 0.02°C/y | 0.04°C/y | 0.06°C/y |
|---|---|---|---|---|---|---|
| Baseline | Ancestral | All | **0%** [0% to 0%] | **+44%** [-11% to +204%] | **+93%** [-13% to +627%] | **+138%** [-13% to +1322%] |
| Evolutionary deviations from baseline | Hot | All | **+10%** [+3% to +35%] | **+12%** [+4% to +43%] | **+19%** [+8% to +109%] | **+24%** [+8% to +204%] |
| | | Development | **-13%** [-40% to -2%] | **-18%** [-84% to -2%] | **-20%** [-150% to -1%] | **-22%** [-247% to -1%] |
| | | Fecundity | **+7%** [+4% to +34%] | **+10%** [+3% to +86%] | **+14%** [+4% to +190%] | **+18%** [+4% to +351%] |
| | | Growth | **+4%** [-2% to +8%] | **+7%** [-1% to +23%] | **+9%** [+2% to +55%] | **+12%** [+3% to +107%] |
| | | Viability | **+8%** [+4% to +39%] | **+9%** [+4% to +64%] | **+10%** [+5% to +93%] | **+12%** [+6% to +121%] |
| | Cold | All | **0%** [-13% to +65%] | **-5%** [-22% to +127%] | **-8%** [-40% to +219%] | **-10%** [-113% to +347%] |
| | | Development | **-1%** [-5% to +8%] | **-1%** [-8% to +8%] | **-2%** [-12% to +12%] | **-2%** [-19% to +23%] |
| | | Fecundity | **-5%** [-13% to +29%] | **-7%** [-34% to +66%] | **-9%** [-70% to +133%] | **-11%** [-147% to +234%] |
| | | Growth | **0%** [-2% to +4%] | **+1%** [-3% to +4%] | **+1%** [-2% to +8%] | **+2%** [0% to +17%] |
| | | Viability | **+3%** [-4% to +32%] | **+2%** [-5% to +49%] | **-1%** [-5% to +63%] | **-3%** [-7% to +66%] |

**Table S3.** Predicted changes in crop damage (per-capita offspring growth rate, g × day$^{-1}$) under different warming scenarios across 20 Californian (USA) sites by the year 2100. The top row shows changes via strictly plastic effects in the ancestral lines (grey), serving as a baseline of ~29% increase in median crop damage rates per °C of warming. The following rows represent additive deviations from this baseline, representing additional fitness changes introduced by evolution (hot = red, cold = blue). Responses have been calculated for the full phenotypes (i.e., by integrating all four evolved reaction norms; trait = all). To investigate trait-specific contributions, responses have also been calculated separately for each trait (i.e., all other reaction norms fixed to their ancestral states). Boxplots show medians (denoted in bold), interquartile ranges, and full ranges (denoted in brackets; arrows denote ranges exceeding the plots' x-limit) across the sites. Evolutionary deviations (colored) from the ancestral baseline (grey) for all warming rates are displayed in the right column.

| | Selection regime | Trait | Warming rate 0°C/y | 0.02°C/y | 0.04°C/y | 0.06°C/y |
|---|---|---|---|---|---|---|
| Baseline | Ancestral | All | **0%** [0% to 0%] | **+42%** [-10% to +202%] | **+92%** [-13% to +609%] | **+134%** [-13% to +1277%] |
| Evolutionary deviations from baseline | Hot | All | **+44%** [+30% to +70%] | **+63%** [+42% to +178%] | **+84%** [+45% to +386%] | **+108%** [+45% to +711%] |
| | | Development | **-8%** [-34% to -2%] | **-11%** [-65% to -2%] | **-12%** [-105% to 0%] | **-12%** [-164% to -1%] |
| | | Fecundity | **+7%** [+4% to +34%] | **+10%** [+4% to +85%] | **+13%** [+4% to +185%] | **+17%** [+4% to +339%] |
| | | Growth | **+28%** [+2% to +52%] | **+41%** [+14% to +149%] | **+55%** [+22% to +353%] | **+72%** [+22% to +666%] |
| | | Viability | **+8%** [+4% to +40%] | **+9%** [+4% to +63%] | **+10%** [+5% to +91%] | **+12%** [+6% to +119%] |
| | Cold | All | **+9%** [-12% to +62%] | **+9%** [-19% to +124%] | **+11%** [-35% to +223%] | **+14%** [-99% to +367%] |
| | | Development | **0%** [-3% to +4%] | **-1%** [-5% to +5%] | **-1%** [-8% to +7%] | **-1%** [-13% to +13%] |
| | | Fecundity | **-5%** [-13% to +29%] | **-6%** [-33% to +65%] | **-9%** [-68% to +130%] | **-11%** [-142% to +227%] |
| | | Growth | **+4%** [-4% to +20%] | **+6%** [-3% to +31%] | **+8%** [+1% to +56%] | **+11%** [+1% to +100%] |
| | | Viability | **+3%** [-4% to +32%] | **+1%** [-5% to +48%] | **-1%** [-5% to +62%] | **-3%** [-7% to +66%] |

**Table S4.** Estimating fitness and crop damage changes using hourly or daily average temperatures (*cf.* Tab. S2 and Tab. S3).

**ORIGINAL SIMULATIONS WITH DIURNAL VARIATION**

| | Selection regime | Trait | Warming rate 0°C/y | 0.02°C/y | 0.04°C/y | 0.06°C/y |
|---|---|---|---|---|---|---|
| Average crop damage rate | Ancestral | All | 0% [0% to 0%] | +42% [-10% to +202%] | +92% [-13% to +609%] | +134% [-13% to +1277%] |
| | Hot | All | +44% [+30% to +70%] | +63% [+42% to +178%] | +84% [+45% to +386%] | +108% [+45% to +711%] |
| | Cold | All | +9% [-12% to +62%] | +9% [-19% to +124%] | +11% [-35% to +223%] | +14% [-99% to +367%] |

| | Selection regime | Trait | 0°C/y | 0.02°C/y | 0.04°C/y | 0.06°C/y |
|---|---|---|---|---|---|---|
| Average fitness (offspring production rate) | Ancestral | All | 0% [0% to 0%] | +44% [-11% to +204%] | +93% [-13% to +627%] | +138% [-13% to +1322%] |
| | Hot | All | +10% [+3% to +35%] | +12% [+4% to +43%] | +19% [+8% to +109%] | +24% [+8% to +204%] |
| | Cold | All | 0% [-13% to +65%] | -5% [-22% to +127%] | -8% [-40% to +219%] | -10% [-113% to +347%] |

**SIMULATIONS WITH DAILY MEANS**

| | Selection regime | Trait | Warming rate 0°C/y | 0.02°C/y | 0.04°C/y | 0.06°C/y |
|---|---|---|---|---|---|---|
| Average crop damage rate | Ancestral | All | 0% [0% to 0%] | +55% [-3% to +339%] | +119% [-7% to +1125%] | +197% [-9% to +2430%] |
| | Hot | All | +46% [-1% to +51%] | +65% [+34% to +201%] | +94% [+44% to +523%] | +122% [+42% to +1149%] |
| | Cold | All | +2% [-16% to +63%] | +4% [-27% to +165%] | +6% [-84% to +356%] | +9% [-183% to +657%] |

| | Selection regime | Trait | 0°C/y | 0.02°C/y | 0.04°C/y | 0.06°C/y |
|---|---|---|---|---|---|---|
| Average $R_0 \times G^{-1}$ | Ancestral | All | 0% [0% to 0%] | +61% [-2% to +344%] | +130% [-7% to +1171%] | +218% [-10% to +2598%] |
| | Hot | All | +6% [-21% to +22%] | +8% [-51% to +75%] | +13% [-42% to +182%] | +20% [+2% to +463%] |
| | Cold | All | -2% [-16% to +70%] | -7% [-29% to +180%] | -13% [-91% to +368%] | -20% [-205% to +645%] |

**Table S5.** Estimating fitness and crop damage changes using different extrinsic mortality rates (adult lifespan / juvenile lifespan; *cf.* Tab. S2 and Tab. S3).

#### ORIGINAL SIMULATIONS WITH INTERMEDIATE EXTRINSIC MORTALITY (AVERAGE LIFESPAN 10/20 DAYS)

| | Selection regime | Trait | Warming rate 0°C/y | 0.02°C/y | 0.04°C/y | 0.06°C/y |
|---|---|---|---|---|---|---|
| Average crop damage rate | Ancestral | All | 0% [0% to 0%] | +42% [-10% to +202%] | +92% [-13% to +609%] | +134% [-13% to +1277%] |
| | Hot | All | +44% [+30% to +70%] | +63% [+42% to +178%] | +84% [+45% to +386%] | +108% [+45% to +711%] |
| | Cold | All | +9% [-12% to +62%] | +9% [-19% to +124%] | +11% [-35% to +223%] | +14% [-99% to +367%] |

| | Selection regime | Trait | 0°C/y | 0.02°C/y | 0.04°C/y | 0.06°C/y |
|---|---|---|---|---|---|---|
| Average fitness (offspring production rate) | Ancestral | All | 0% [0% to 0%] | +44% [-11% to +204%] | +93% [-13% to +627%] | +138% [-13% to +1322%] |
| | Hot | All | +10% [+3% to +35%] | +12% [+4% to +43%] | +19% [+8% to +109%] | +24% [+8% to +204%] |
| | Cold | All | 0% [-13% to +65%] | -5% [-22% to +127%] | -8% [-40% to +219%] | -10% [-113% to +347%] |

#### SIMULATIONS WITH HIGH EXTRINSIC MORTALITY (AVERAGE LIFESPAN 5/10 DAYS)

| | Selection regime | Trait | Warming rate 0°C/y | 0.02°C/y | 0.04°C/y | 0.06°C/y |
|---|---|---|---|---|---|---|
| Average crop damage rate | Ancestral | All | 0% [0% to 0%] | +66% [-14% to +445%] | +141% [-18% to +1898%] | +216% [-19% to +4906%] |
| | Hot | All | +31% [-2% to +66%] | +57% [+32% to +172%] | +87% [+34% to +635%] | +128% [+33% to +1723%] |
| | Cold | All | +5% [-13% to +67%] | +6% [-28% to +243%] | +6% [-114% to +652%] | +9% [-474% to +1352%] |

| | Selection regime | Trait | 0°C/y | 0.02°C/y | 0.04°C/y | 0.06°C/y |
|---|---|---|---|---|---|---|
| Average fitness (offspring production rate) | Ancestral | All | 0% [0% to 0%] | +68% [-14% to +454%] | +150% [-18% to +1974%] | +223% [-19% to +5210%] |
| | Hot | All | -1% [-18% to +36%] | +1% [-82% to +40%] | +4% [-193% to +45%] | +16% [-335% to +193%] |
| | Cold | All | -4% [-13% to +70%] | -9% [-29% to +245%] | -12% [-130% to +632%] | -14% [-526% to +1250%] |

#### SIMULATIONS WITH LOW EXTRINSIC MORTALITY (AVERAGE LIFESPAN 20/40 DAYS)

| | Selection regime | Trait | Warming rate 0°C/y | 0.02°C/y | 0.04°C/y | 0.06°C/y |
|---|---|---|---|---|---|---|
| Average crop damage rate | Ancestral | All | 0% [0% to 0%] | +33% [-7% to +145%] | +68% [-9% to +325%] | +100% [-7% to +625%] |
| | Hot | All | +53% [+38% to +92%] | +67% [+46% to +177%] | +83% [+52% to +304%] | +102% [+51% to +474%] |
| | Cold | All | +11% [-11% to +58%] | +11% [-16% to +89%] | +13% [-23% to +134%] | +15% [-38% to +195%] |

| | Selection regime | Trait | 0°C/y | 0.02°C/y | 0.04°C/y | 0.06°C/y |
|---|---|---|---|---|---|---|
| Average fitness (offspring production rate) | Ancestral | All | 0% [0% to 0%] | +33% [-8% to +147%] | +72% [-9% to +332%] | +102% [-8% to +644%] |
| | Hot | All | +17% [+8% to +43%] | +19% [+10% to +63%] | +24% [+13% to +109%] | +30% [+13% to +167%] |
| | Cold | All | +2% [-12% to +62%] | -2% [-17% to +92%] | -4% [-25% to +135%] | -6% [-42% to +190%] |
